# BOLD and EEG Signal Variability at Rest Differently Relate to Aging in the Human Brain

**DOI:** 10.1101/646273

**Authors:** D. Kumral, F. Şansal, E. Cesnaite, K. Mahjoory, E. Al, M. Gaebler, V. V. Nikulin, A. Villringer

## Abstract

Variability of neural activity is regarded as a crucial feature of healthy brain function, and several neuroimaging approaches have been employed to assess it noninvasively. Studies on the variability of both evoked brain response and spontaneous brain signals have shown remarkable changes with aging but it is unclear if the different measures of brain signal variability – identified with either hemodynamic or electrophysiological methods – reflect the same underlying physiology. In this study, we aimed to explore age differences of spontaneous brain signal variability with two different imaging modalities (EEG, fMRI) in healthy younger (25±3 years, N=135) and older (67±4 years, N=54) adults. Consistent with the previous studies, we found lower blood oxygenation level dependent (BOLD) variability in the older subjects as well as less signal variability in the amplitude of low-frequency oscillations (1–12 Hz), measured in source space. These age-related reductions were mostly observed in the areas that overlap with the default mode network. Moreover, age-related increases of variability in the amplitude of beta-band frequency EEG oscillations (15–25 Hz) were seen predominantly in temporal brain regions. There were significant sex differences in EEG signal variability in various brain regions while no significant sex differences were observed in BOLD signal variability. Bivariate and multivariate correlation analyses revealed no significant associations between EEG- and fMRI-based variability measures. In summary, we show that both BOLD and EEG signal variability reflect aging-related processes but are likely to be dominated by different physiological origins, which relate differentially to age and sex.

## 1. Introduction

Functional neuroimaging methods such as fMRI, PET, fNIRS, EEG, or MEG have allowed the non-invasive assessment of functional changes in the aging human brain (Cabeza, 2001; Cabeza et al., 2018). Most previous functional neuroimaging studies on aging have employed a task-based design (Grady, 2012) and in their data analysis the central tendency has typically been assumed to be the most representative value in a distribution (e.g., mean) (Speelman and McGann, 2013) or the “signal” within distributional “noise”. In recent years, also the variability of brain activation in task-dependent and task-independent measurements (as spontaneous variations of background activity) has been shown to provide relevant information about the brain’s functional state (Garrett et al., 2013b; Grady and Garrett, 2018; Nomi et al., 2017). These studies primarily measured the blood oxygen level dependent (BOLD) signal using fMRI. For example, it has been demonstrated that the variance of the task-evoked BOLD response was differentially related to aging as well as cognitive performance (Armbruster-Genc et al., 2016; Garrett et al., 2013a). Similarly, spontaneous signal variability in resting state fMRI (rsfMRI) has been found to decrease with age (Grady and Garrett, 2018; Nomi et al., 2017), in individuals with stroke (Kielar et al., 2016), and 22q11.2 deletion syndrome (Zöller et al., 2017). An increase of fMRI variability has been shown to occur in inflammation induced state-anxiety (Labrenz et al., 2018) and to parallel symptom severity in Attention Deficit Hyperactivity Disorder (Nomi et al., 2018). From these studies, it was concluded that changes in BOLD signal variability might serve as an index for alterations in neural processing and cognitive flexibility (Grady and Garrett, 2014).

The conclusions of aforementioned studies imply that BOLD signal variability is mainly determined by *neuronal* variability. To a large extent, this is based on the premise that BOLD is related to neuronal activity: The evoked BOLD signal in task-based fMRI reflects the decrease of the deoxyhemoglobin concentration to changes in local brain activity, which is determined by vascular (blood velocity and volume: “neurovascular coupling”) and metabolic (oxygen consumption: “neurometabolic coupling”) factors (Logothetis and Wandell, 2004; Villringer and Dirnagl, 1995). The BOLD signal is therefore only an indirect measure of neural activity (Logothetis, 2008). For the variability of task-evoked BOLD signal and for spontaneous variations of the BOLD signal, in principle, the same considerations apply regarding their relationship to underlying neural processes (Murayama et al., 2010). However, since in rsfMRI there is no explicit external trigger for evoked brain activity to which time-locked averaging could be applied, the time course of rsfMRI signals is potentially more susceptible to contributions of “physiological noise”, such as cardiac and respiratory signals (Birn et al., 2008; Chang et al., 2009), but also spontaneous fluctuations of vascular tone, which is found even in isolated arterial vessels (Failla et al., 1999; Hudetz et al., 1998; Wang et al., 2006). In the same vein, the variability of task-evoked fMRI is not necessarily reflecting only the variability of evoked neuronal activity, as it may also – at least partly – reflect the variability of the spontaneous background signal on which a constant evoked response is superimposed.

In aging, non-neuronal signal fluctuations may also introduce spurious common variance across the rsfMRI time series (Caballero-Gaudes and Reynolds, 2017), thus confounding estimates of “neural” brain signal variability. Previous evidence suggests that the relationship between neuronal activity and the vascular response is attenuated with age – and so is, as a consequence, the BOLD signal (for review see D’Esposito et al., 2003). For instance, aging has been associated with altered cerebrovascular ultrastructure, reduced elasticity of vessels, and atherosclerosis (Farkas and Luiten, 2001) but also with a decrease in resting cerebral blood flow (CBF), cerebral metabolic rate of oxygen (CMRO_2_), and cerebrovascular reactivity (CVR) (Liu et al., 2013). Taken together, age-related changes in BOLD signal or BOLD signal variability are related to a mixture of alterations in non-neural spontaneous fluctuations of vascular signals, neural activity, neurovascular coupling, and/or neurometabolic coupling (D’Esposito et al., 2003; Geerligs et al., 2017; Tsvetanov et al., 2015).

While BOLD fMRI signal and specifically variance measures based on fMRI are only partially and indirectly related to neural activity (Liu, 2013; Logothetis, 2008), electrophysiological methods such as EEG can provide a more direct assessment of neural activity with a higher temporal but poorer spatial resolution (Cohen, 2017). EEG measures neuronal currents resulting from the synchronization of dendritic postsynaptic potentials across the neural population; the cerebral EEG rhythms thereby reflect the underlying brain neural network activity (Steriade, 2006). Resting state (rs)EEG is characterized by spontaneous oscillations (“brain rhythms”) at different frequencies. Previously, the mean amplitude of low-frequency bands (e.g., delta and/or theta, 1-7 Hz) has been shown to correlate negatively with age (Vlahou et al., 2015), while higher-frequency bands (e.g., beta, 15-25 Hz) show the reverse pattern (Rossiter et al., 2014). However, less is known about the within-subject variability of EEG measures and their association with aging. Several studies have addressed the variability in the spectral amplitudes of different frequency bands using variance (Hawkes and Prescott, 1973; Oken and Chiappa, 1988), coefficient of variation (Burgess and Gruzelier, 1993; Maltez et al., 2004), and complexity (Fernández et al., 2012; Sleimen-Malkoun et al., 2015). For instance, reductions of the complexity in rsEEG signal have been found not only in healthy aging (Yang et al., 2013; Zappasodi et al., 2015) but also in age-related pathologies such as mild cognitive impairment (McBride et al., 2014) and Alzheimer’s disease (Smits et al., 2016). Accordingly, it has been suggested that irregular (e.g., variable) systems indicate a normal and healthy state (more integrated information) while highly regular systems often mark dysfunction or disease (Lipsitz and Goldberger, 1992; Vaillancourt and Newell, 2002).

The different methodological approaches, fMRI based “vascular” approaches on the one hand and electrophysiological methods such as EEG and MEG, on the other hand, indicate alterations of brain signal variability with aging. However, it remains unclear whether these different measures of brain variability at rest reflect the same underlying physiological changes. Evidently, there are some correlations between the two signal sources (for a review see, Jorge et al., 2014; Ritter and Villringer, 2006). For instance, in task-based EEG-fMRI simultaneous recordings, a relationship between BOLD responses and amplitude of evoked potentials has been demonstrated (e.g., Ritter et al., 2009; Seaquist et al., 2007), while in resting state EEG-fMRI studies, a negative association between spontaneous modulations of alpha rhythm and BOLD signal has also been established (e.g., Chang et al., 2013; Goldman et al., 2002; Gonçalves et al., 2006; Moosmann et al., 2003). Further, differential correlation patterns have been noted for the various rhythms of different frequencies in EEG/MEG and the fMRI signal, such that low-frequency oscillations show a negative (Deligianni et al., 2014; Mantini et al., 2007; Meyer et al., 2013), while higher frequencies oscillations demonstrate a positive correlation with the BOLD signal (Niessing et al., 2005; Scheeringa et al., 2011).

Regarding the known age-related changes in BOLD and EEG signal variability, respectively, the question arises whether these alterations are dominated by joint signal sources of fMRI and EEG or by – potentially different – signal contributions that relate to each of these two methods. Given the – potentially large – non-neuronal signal contribution, this issue is particularly relevant for rsfMRI studies. Here, we addressed this question by analyzing rsfMRI and EEG measures of variability in healthy younger and older subjects. To our knowledge, the only study that compared variability in a “vascular” imaging method (rsfMRI) and an electrophysiological method (rsMEG at the sensor space) concluded that the effects of aging on BOLD signal variability were mainly driven by vascular factors (e.g., heart rate variability) and not well-explained by the changes in neural variability (Tsvetanov et al., 2015). The main aims of the present study were to explore i) age differences of brain signal variability measures, as well as to investigate ii) how neural variability derived from rsEEG related to the analogous parameters of BOLD signal variability derived from rsfMRI. We used rsfMRI and rsEEG from the “Leipzig Study for Mind-Body-Emotion Interactions” (Babayan et al., 2019). As an explanatory analysis, we further investigated sex-related differences of brain signal variability measures. To measure brain signal variability, we calculated the standard deviation (SD) of both the BOLD signal and of the amplitude envelope of the filtered rsEEG time series for a number of standard frequency bands at the source space. We hypothesized that brain signal variability would generally decrease with aging. In addition, based on the premise that BOLD fMRI signal variability reflects *neural* variability as measured by rsEEG, we expected that the corresponding changes in both signal modalities would demonstrate moderate to strong similarity in their spatial distribution. Given the confounding effects of vascular factors during aging on the fMRI signal (D’Esposito et al., 2003; Liu, 2013; Thompson, 2018), we further expected to find the relationship between BOLD and EEG signal variability to be stronger in younger than older adults.

## 2. Method

### 2.1. Participants

The data of the “Leipzig Study for Mind-Body-Emotion Interactions” (LEMON; Babayan et al., 2019) comprised 227 subjects in two age groups (younger: 20-35, older: 59-77). Only participants who did not report any neurological disorders, head injury, alcohol or other substance abuse, hypertension, pregnancy, claustrophobia, chemotherapy and malignant diseases, current and/or previous psychiatric disease or any medication affecting the cardiovascular and/or central nervous system in a telephone pre-screening were invited to the laboratory. The study protocol conformed to the Declaration of Helsinki and was approved by the ethics committee at the medical faculty of the University of Leipzig (reference number 154/13-ff).

RsEEG recordings were available for 216 subjects who completed the full study protocol. We excluded data from subjects that had missing event information (N=1), different sampling rate (N=3), mismatching header files or insufficient data quality (N=9). Based on the rsfMRI quality assessment, we further excluded data from subjects with faulty preprocessing (N=7), ghost artefacts (N=2), incomplete data (N=1), or excessive head motion (N=3) (criterion: mean framewise displacement (FD) ≤ 0.5 mm; Power et al., 2012) (Supplementary Figure 1). The final sample included 135 younger (*M* = 25.10 ± 3.70 years, 42 females) and 54 older subjects (*M* = 67.15 ± 4.52 years, 27 females).

### 2.1. fMRI Acquisition

Brain imaging was performed on a 3T Siemens Magnetom Verio MR scanner (Siemens Medical Systems, Erlangen, Germany) with a standard 32-channel head coil. The participants were instructed to keep their eyes open and not fall asleep while looking at a low-contrast (light grey on dark grey background) fixation cross.

The structural image was recorded using an MP2RAGE sequence (Marques et al., 2010) with the following parameters: TI 1 = 700 ms, TI 2 = 2500 ms, TR = 5000 ms, TE = 2.92 ms, FA 1 = 4°, FA 2 = 5°, band width = 240 Hz/pixel, FOV = 256 × 240 × 176 mm^3^, voxel size = 1 × 1 × 1 mm^3^. The functional images were acquired using a T2*-weighted multiband EPI sequence with the following parameters: TR = 1400 ms, TE = 30 ms, FA= 69°, FOV = 202 mm, imaging matrix=88 × 88, 64 slices with voxel size = 2.3 × 2.3 × 2.3 mm^3^, slice thickness = 2.3 mm, echo spacing = 0.67 ms, bandwidth=1776 Hz/Px, partial fourier 7/8, no pre-scan normalization, multiband acceleration factor = 4, 657 volumes, duration = 15 min 30 s. A gradient echo field map with the sample geometry was used for distortion correction (TR = 680 ms, TE 1 = 5.19 ms, TE 2 = 7.65 ms).

### 2.2. fMRI Preprocessing

Preprocessing was implemented in Nipype (Gorgolewski et al., 2011), incorporating tools from FreeSurfer (Fischl, 2012), FSL (Jenkinson et al., 2012), AFNI (Cox, 1996), ANTs (Avants et al., 2011), CBS Tools (Bazin et al., 2014), and Nitime (Rokem et al., 2009). The pipeline comprised the following steps: (I) discarding the first five EPI volumes to allow for signal equilibration and steady state, (II) 3D motion correction (FSL mcflirt), (III) distortion correction (FSL fugue), (IV) rigid body coregistration of functional scans to the individual T1-weighted image (Freesurfer bbregister), (V) denoising including removal of 24 motion parameters (CPAC, Friston et al., 1996), motion, signal intensity spikes (Nipype rapidart), physiological noise in white matter and cerebrospinal fluid (CSF) (CompCor; Behzadi et al., 2007), together with linear and quadratic signal trends, (VI) band-pass filtering between 0.01-0.1 Hz (FSL fslmaths), (VII) spatial normalization to MNI152 (Montreal Neurological Institute) standard space (2 mm isotropic) via transformation parameters derived during structural preprocessing (ANTS). (VIII) The data were then spatially smoothed with a 6-mm full-width half-maximum (FWHM) Gaussian kernel (FSL fslmaths). Additionally, we calculated total intracranial volume (TIV) of each subject using the Computational Anatomy Toolbox (CAT12: http://dbm.neuro.uni-jena.de/cat/) running on Matlab 9.3 (Mathworks, Natick, MA, USA) and used it as a covariate for further statistical analyses (Malone et al., 2015).

#### BOLD Signal Variability (SD_BOLD_)

Standard deviation (SD) quantifies the amount of variation or dispersion in a set of values (Garrett et al., 2015; Grady and Garrett, 2018). Higher SD in rsfMRI signal indicates greater intensity of signal fluctuation or an increased level of activation in a given area (Garrett et al., 2011). We first calculated SD_BOLD_ across the whole time series for each voxel and then within 96 boundaries of preselected atlas-based regions of interests (ROIs) based on the Harvard-Oxford cortical atlas (Desikan et al., 2006). The main steps of deriving brain signal variability (SD_BOLD_) from the preprocessed fMRI signal are shown in Figure 1.

**Figure 1.**
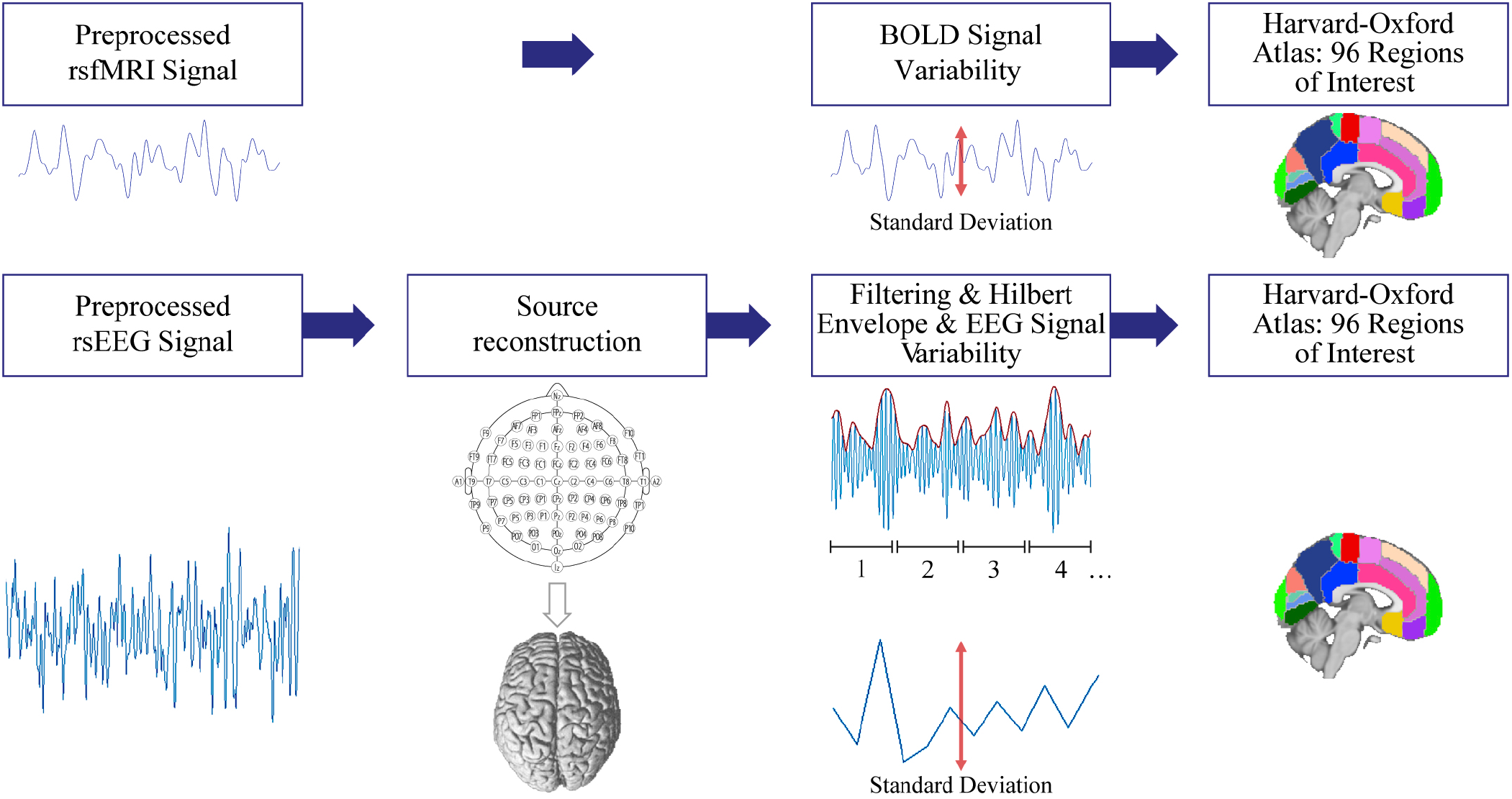
Main steps of deriving brain signal variability from the preprocessed resting state fMRI and EEG signal. We calculated the standard deviation of the blood oxygen level dependent (BOLD) signal and of the coarse-grained amplitude envelope of the rsEEG time series for a number of standard frequency bands at the source space. Each sample of coarse-grained amplitude envelope of the rsEEG (represented in numbers) is generated by averaging the samples in non-overlapping windows of length 0.5 s.

The reproducible workflows containing fMRI preprocessing details can be found here: https://github.com/NeuroanatomyAndConnectivity/pipelines/releases/tag/v2.0.

### 2.3. EEG Recordings

Sixteen minutes of rsEEG were acquired on a separate day with BrainAmp MR-plus amplifiers using 61 ActiCAP electrodes (both Brain Products, Germany) attached according to the international standard 10-20 localization system (Jurcak et al., 2007) with FCz (fronto-central or cephalic electrode) as the reference. The ground electrode was located at the sternum. Electrode impedance was kept below 5 kΩ. Continuous EEG activity was digitized at a sampling rate of 2500 Hz and band–pass filtered online between 0.015 Hz and 1 kHz.

The experimental session was divided into 16 blocks, each lasting 60 s, with two conditions interleaved, eyes closed (EC) and eyes open (EO), starting with the EC condition. Changes between blocks were announced with the software Presentation (v16.5, Neurobehavioral Systems Inc., USA). Participants were asked to sit comfortably in a chair in a dimly illuminated, sound-shielded Faraday recording room. During the EO periods, participants were instructed to stay awake while fixating on a black cross presented on a white background. To maximize comparability, only EEG data from the EO condition were analyzed, since rsfMRI data were collected only in the EO condition.

### 2.4. EEG Data Analysis

EEG processing and analyses were performed with custom Matlab (The MathWorks, Inc, Natick, Massachusetts, USA) scripts using functions from the EEGLAB environment (version 14.1.1b; Delorme and Makeig, 2004). The continuous EEG data were down-sampled to 250 Hz, band-pass filtered within 1–45 Hz (4^th^ order back and forth Butterworth filter) and split into EO and EC conditions. Segments contaminated by large artefacts due to facial muscle tensions and gross movements were removed following visual inspection, resulting in a rejection of on average 6.6% of the recorded data. Rare occasions of artifactual channels were excluded from the analysis. The dimensionality of the data was reduced using principal component analysis (PCA) by selecting at least 30 principal components explaining 95% of the total variance. Next, using independent component analysis (Infomax; Bell and Sejnowski, 1995), the confounding sources e.g. eye-movements, eye-blinks, muscle activity, and residual ballistocardiographic artefacts were rejected from the data.

### 2.5. EEG Source Reconstruction

Before conducting source reconstruction, preprocessed EEG signals were re-referenced to a common average. We incorporated a standard highly detailed finite element method (FEM) volume conduction model as described by Huang et al. (2016). The geometry of the FEM model was based on an extended MNI/ICBM152 (International Consortium for Brain Mapping) standard anatomy, where the source space constrained to cortical surface and parceled to 96 ROIs based on the Harvard-Oxford atlas (Desikan et al., 2006). We used eLORETA (exact low resolution brain electromagnetic tomography) as implemented in as implemented in as implemented in the M/EEG Toolbox of Hamburg (METH; Haufe and Ewald, 2016; Pascual-Marqui, 2007) to compute the cortical electrical distribution from the scalp EEG recordings. The leadfield matrix was calculated between 1804 points located on the cortical surface to the 61 scalp electrodes. We filtered into several frequency bands, associated with brain oscillations: delta (1–3 Hz), theta (4–8 Hz), alpha (8–12 Hz), and beta (15–25 Hz). Following the singular value decomposition (SVD) of each voxel’s three-dimensional time course, the dominant orientation of the source signal was identified by preserving the first SVD component. The amplitude envelope of filtered oscillations was extracted using the Hilbert transform (Rosenblum et al., 2001). Next, we applied temporal coarse graining by averaging data points in non-overlapping windows of length 0.5 s (Figure 1).

#### EEG Variability (SD_EEG_)

We calculated the SD of amplitude envelope of band-pass filtered oscillations on the coarse-grained signal. RsEEG signal variability (SD_EEG_) was obtained for different frequency bands (SD_DELTA_, SD_THETA_, SD_ALPHA_, SD_BETA_) in each of 96 ROIs. Further, in our study we investigated variability in the amplitude of oscillatory signals from one time segment to the other. If amplitude (or power) of each signal stays the same, the variability (SD) in the amplitude of such segments will be zero. Therefore, the average amplitude of a signal is not indicative of its variability. Although amplitude and its standard deviation mathematically are different, they can show some correlation due to size effects (Immer, 1937).

Main steps toward deriving brain signal variability from the preprocessed EEG signal are shown in Figure 1. The raw and preprocessed fMRI and EEG data samples can be found at https://ftp.gwdg.de/pub/misc/MPI-Leipzig_Mind-Brain-Body-LEMON/

### 2.6. Statistical Analyses

#### Mean SD_BOLD_ and SD_EEG_

For the topographic information (based on ROIs), the mean BOLD and EEG variability were calculated by I) log-transforming the SD values, II) averaging separately for younger and older subjects, and III) then back-transforming the values (McDonald, 2014).

#### Age and Sex Effects

A series of non-parametric analyses of covariance (ANCOVAs, type III) were applied to brain signal variability values in each 96 ROIs for SD_BOLD_ and SD_EEG_ using age group and sex as variables of interest, adjusting for TIV and mean FD. The significance level was controlled for using false discovery rate (FDR) correction according to Benjamini and Hochberg (1995). Significant group differences were further examined by Tukey HSD post-hoc comparisons. The signal variability values were log-transformed to normalize SD_BOLD_ and SD_EEG_ before further analyses (assessed by Lilliefors tests at a significance threshold of 0.05). All analyses were performed using the *aovp* function in the *lmperm* package (Wheeler, 2016) as implemented in R (R core team, 2018).

#### SD_BOLD_ – SD_EEG_ Correlation

To investigate the association between each ROI of SD_BOLD_ and SD_EEG_, we used pairwise Spearman’s rank correlation separately for younger and older subjects, corrected for FDR (96 ROIs). We further applied sparse canonical correlation analysis (CCA) to show that the relationship between SD_BOLD_ and SD_EEG_ is not missed when only mass bivariate correlations are used. CCA is a multivariate method to find the independent linear combinations of variables such that the correlation between variables is maximized (Witten et al., 2009). The sparse CCA criterion is obtained by adding a Lasso Penalty function (*l*_1_), which performs continuous shrinkage and automatic variable selection and can solve statistical problems such as multicollinearity and overfitting (Tibshirani, 2011). We used *l*_1_ penalty as the regularization function to obtain sparse coefficients, that is, the canonical vectors (i.e., translating from full variables to a data matrix’s low-rank components of variation) will contain exactly zero elements. Sparse CCA was performed using the R package PMA (Penalized Multivariate Analysis; Witten et al., 2009; http://cran.r-project.org/web/packages/PMA/). In our analyses, the significance of the correlation was estimated using the permutation approach (N=1000) as implemented in the CCA.permute function in R (p_perm_<0.05).

#### Cognition

The Trail Making Test (TMT) is a cognitive test measuring executive function, including processing speed and mental flexibility (Reitan, 1955; Reitan and Wolfson, 1995). In the first part of the test (TMT-A) the targets are all numbers, while in the second part (TMT-B), participants need to alternate between numbers and letters. In both TMT-A and B, the time to complete the task quantifies the performance, and lower scores indicate better performance. Based on the previous literature, we focused on SD_BOLD_, SD_DELTA_, and SD_THETA_ (Vlahou et al., 2015) and selected different ROIs from two research papers about the neural correlates of the TMT: Zakzanis et al., (2005) and Jacobson et al., (2011) (Table 1). To reduce the number of multiple comparisons (Nguyen and Holmes, 2019), these ROIs were decomposed into singular values using the *prcomp* function belonging to *factoextra* package (R core team, 2018), which performs SVD on the centered values. As a criterion, the minimum total variance explained over 70% was selected (Jollife and Cadima, 2016). This resulted in three principle components (PC) in SD_BOLD_ (52.82%, 10.34%, and 7%), two PCs in SD_DELTA_ (67.37%, 10.95%), and one PC in SD_THETA_ (75.63%). We also ran multiple linear regression using task completion time in TMT-A and TMT-B as the dependent variables with the PC scores (for SD_BOLD_, SD_DELTA_, and SD_THETA_) and their interaction with continuous age as independent variables. Since the residuals from the regression models fitted to the data were not normally distributed, the TMT values were log-transformed prior to the final analyses. These tests were conducted using the *lmp* function in *lmperm* package implemented in R (R core team, 2018).

**Table 1.**
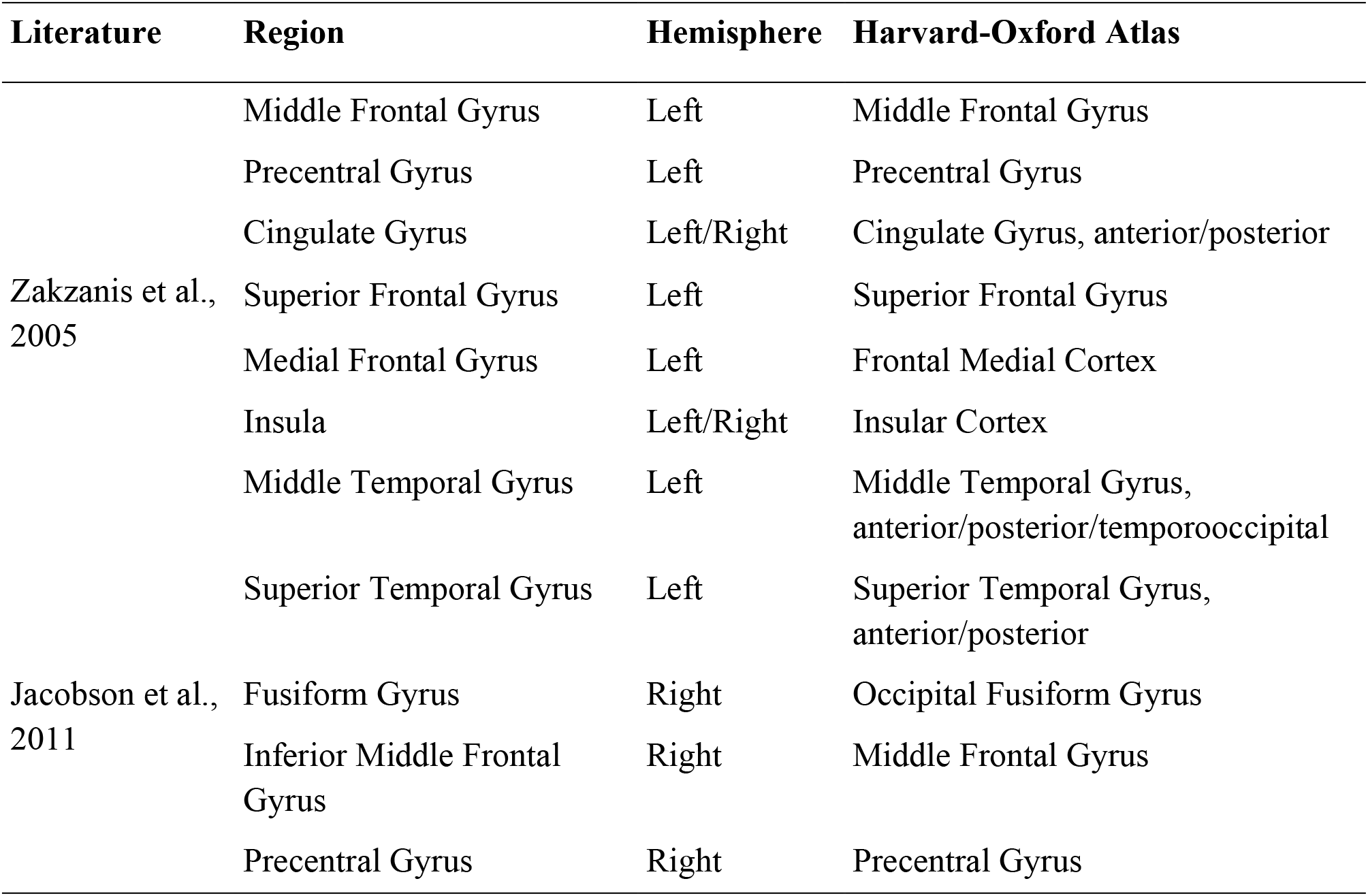
Selected region of interests (ROIs) derived from the previous fMRI literature, and their corresponding ROIs in Harvard-Oxford atlas to investigate the age-dependent relationship between TMT and SD_BOLD_ or SD_EEG_.

## 3. Results

### Mean SD_BOLD_ and SD_EEG_

The topographic distribution of SD_BOLD_ in younger adults revealed the largest brain signal variability values in fronto-temporal regions while in older adults it was in the frontal and occipital areas. Further, we found strongest variability across younger subjects in occipito-temporal regions for SD_DELTA_, SD_THETA_, SD_ALPHA_, and in medial frontal brain regions for SD_BETA_, while older adults showed strongest brain signal variability in the fronto-central brain regions for SD_DELTA_, in parietal-central brain regions for SD_THETA_, SD_ALPHA_, and in medial frontal brain regions for SD_BETA_. The details of the mean values of SD_BOLD_ and SD_EEG_ across age groups and their topographic distributions are given in Supplementary Table 1, Supplementary Figure 2 and 3, and are also available at Neurovault (https://neurovault.org/collections/WWOKVUDV/).

### Age and Sex Effects

The nonparametric ANCOVAs with SD_BOLD_ as dependent variable demonstrated that there was a significant main effect of age group in 72 ROIs in frontal, temporal, and occipital brain regions (F-values: 13.32–61.14; Figure 2). However, there was no significant main effect of sex on SD_BOLD_ and no significant interaction between age group and sex (all p_FDR_>0.05). Tukey HSD post-hoc analyses showed that older subjects had decreased SD_BOLD_ compared to younger adults which were presented in both sexes (n_ROI_=35). The nonparametric ANCOVAs with SD_EEG_ as dependent variable showed significant main effects of age group in all frequency bands: SD_DELTA_ in 14 ROIs in occipital (F-values: 12.57– 20.94), SD_THETA_ in 16 ROIs in frontal and parietal (F-values: 13.16–40.30), SD_ALPHA_ in 20 ROIs in occipital (F-values: 12.69–20.12), and SD_BETA_ in 19 ROIs in central and temporal brain regions (F-values: 12.50–21.61), as shown in Figure 2. There were also significant main effects of sex in all frequency bands: SD_DELTA_ in 21 ROIs in temporal and occipital (F-values: 13.24–26.63), SD_THETA_ in 74 ROIs in frontal, occipital, and temporal (F-values: 12.68–30.06), SD_ALPHA_ in 4 ROIs in frontal (F-values: 12.88–16.51), and SD_BETA_ in 69 ROIs in temporal, occipital, and central brain regions (F-values: 12.54–35.72), as shown in Figure 3. No significant interaction effects between age group and sex on SD_EEG_ were observed in any frequency band (p_FDR_>0.05). Tukey HSD post-hoc analyses on SD_EEG_ showed that older subjects had less brain signal variability, which was present in both sexes for SD_DELTA_ (n_ROI_=12), SD_THETA_ (n_ROI_=10), and SD_ALPHA_ (n_ROI_=11). Additionally, older adults showed higher SD_BETA_, driven by female subjects (n_ROI_=15). With regard to sex differences, post-hoc analyses showed that females had higher SD_DELTA_, SD_THETA_, SD_ALPHA_, and SD_BETA_ than males. Sex differences in SD_DELTA_ (n_ROI_=13) and SD_THETA_ (n_ROI_=54) were mostly pronounced in younger adults, while the effect of sex in SD_BETA_ (n_ROI_=21) were mainly presented in older adults (p<0.05). The graphical distribution of the F-values for the significant effects of age group or sex for each ROIs are shown in Supplementary Figure 4. Additional information of SD_BOLD_ and SD_EEG_ for each frequency band and for each of the 96 ROIs, split up by age group and sex, are presented in the Supplementary Tables 2-6.

**Figure 2.**
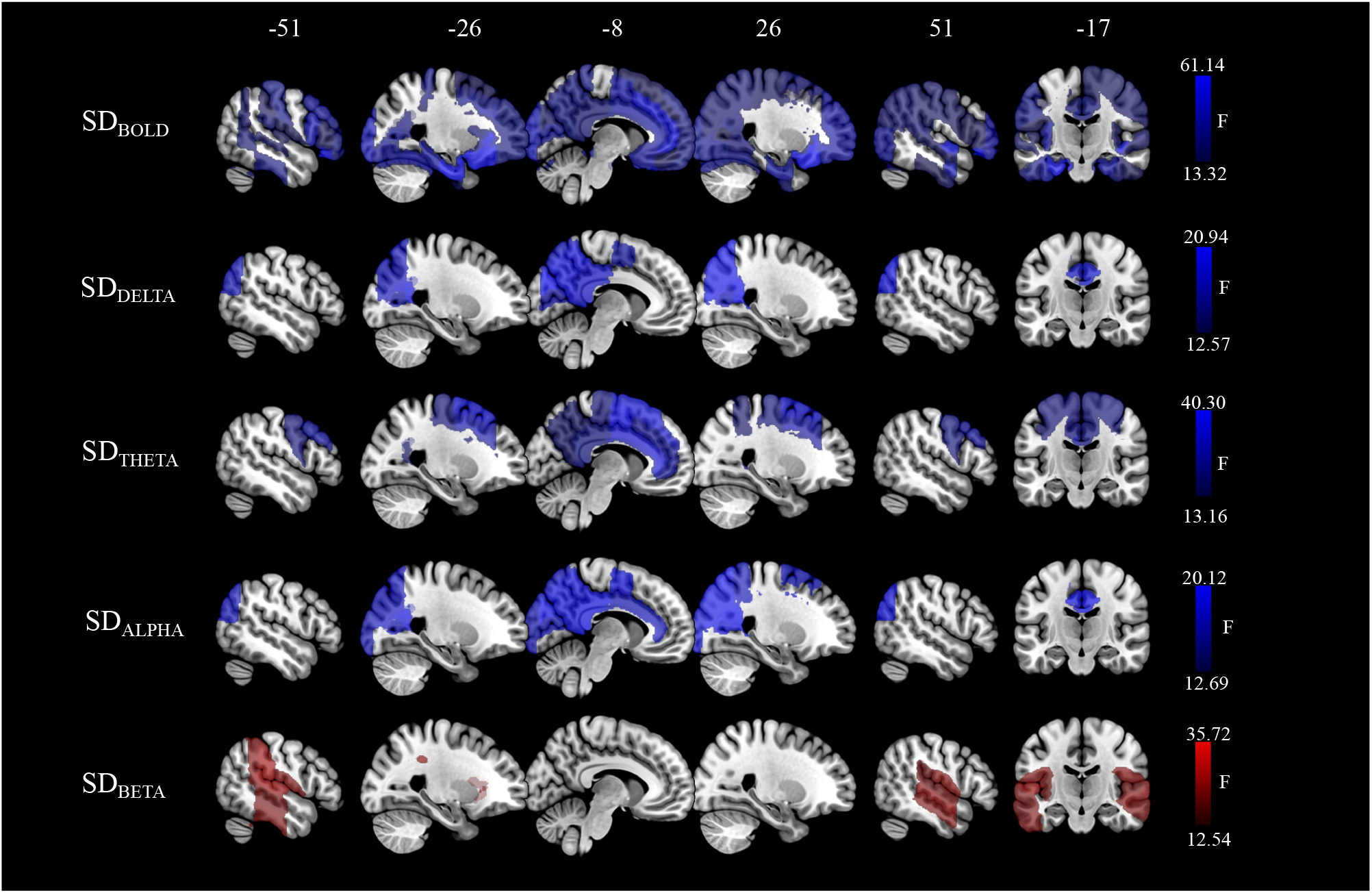
Spatial maps of significant age group differences in SD_BOLD_ and SD_EEG_. We calculated the standard deviation (SD) of the blood oxygen level dependent (BOLD) signal and of the coarse-grained amplitude envelope of the rsEEG time series for the delta (1–3 Hz), theta (4–8 Hz), alpha (8–12 Hz), and beta (15–25 Hz) frequency bands at the source space. Statistical significance was determined using nonparametric ANCOVAs corrected for multiple comparisons by false discovery rates (FDR; Benjamini and Hochberg, 1995). Blue color indicates areas where brain signal variability was lower in older than in younger adults, while red color indicates the opposite.

**Figure 3.**
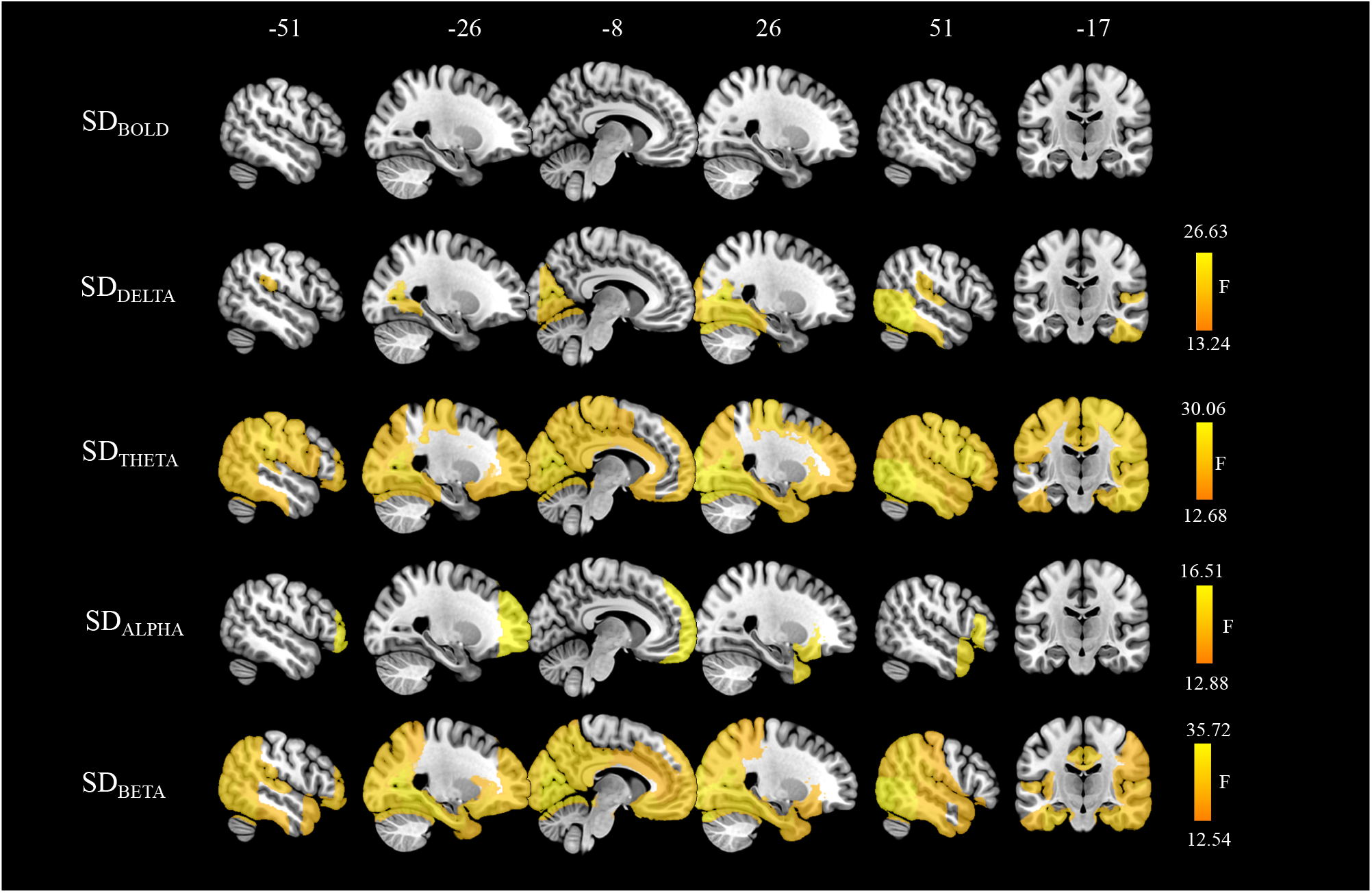
Spatial maps of significant sex differences in SD_BOLD_ and SD_EEG_. We calculated the standard deviation (SD) of the blood oxygen level dependent (BOLD) signal and of the coarse-grained amplitude envelope of the rsEEG time series for the delta (1– 3 Hz), theta (4–8 Hz), alpha (8–12 Hz), and beta (15–25 Hz) frequency bands at the source space. Statistical significance was determined using nonparametric ANCOVAs corrected for multiple comparisons by false discovery rates (FDR; Benjamini and Hochberg, 1995). Yellow color indicates areas where brain signal variability was higher in female subjects as compared to male subjects in EEG.

### SD_BOLD_ – SD_EEG_ Correlation

The correlation coefficient of pairwise associations for 96 ROIs of SD_BOLD_ with SD_DELTA_, SD_THETA_, SD_ALPHA_, and SD_BETA_ ranged in younger adults from rho=-0.200 to rho=0.223 (Supplementary Table 7) and in older adults from rho=0.386 to rho=0.349 (Supplementary Table 8). None of the pairwise associations between SD_BOLD_ and SD_EEG_ remained significant after the correction for multiple comparison corrections. Confirmatory multivariate sparse CCA further showed that correlations between SD_BOLD_ and SD_EEG_ across all subjects were rather low, highly sparse, and non-significant (SD_DELTA_; r=0.145, p_perm_ =0.750, *l*_1_=0.367; SD_THETA_; r=0.143, p_perm_=0.713 *l*_1_=0.7; SD_ALPHA_; r=0.153, p_perm_=0.528, *l*_1_=0.1; SD_BETA_; r=0. 232, p_perm_=0.096, *l*_1_=0.633).

### Cognition

There was a significant interaction between age and SD_BOLD_ in PC2 on the TMT-B performance (adjusted R^2^ = 0.395, *F*(7,181) = 18.60, p <.001, interaction: β= −0.002, p = 0.027). For older, but not younger participants, stronger SD_BOLD_ was associated with faster completion time in PC2, driven mainly by the left temporal gyrus as well as the left anterior and posterior cingulate cortex (Figure 5). The regression analyses in SD_DELTA_ and SD_THETA_ did not show a significant association between cognition and brain signal variability measures. The contributions of selected ROIs to the PCs resulted from SVD analyses can be found in Supplementary Table 9. The complete multiple linear regression results can be found in Supplementary Table 10.

**Figure 4.**
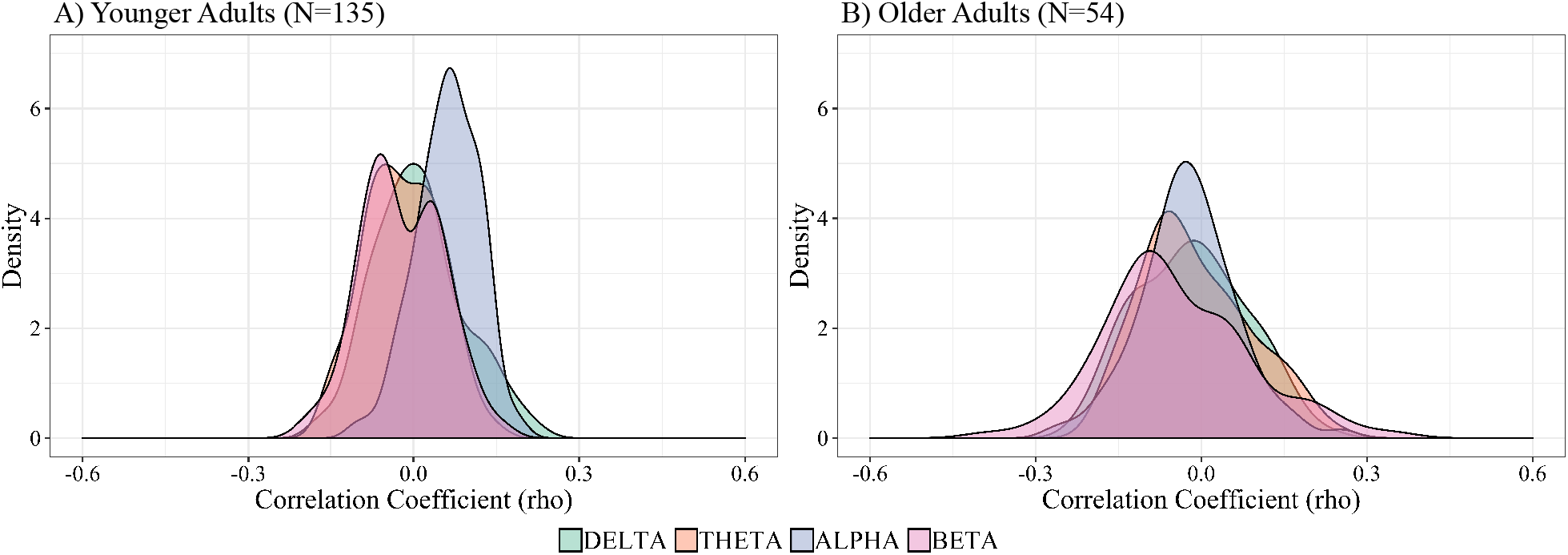
Distribution of correlation coefficients (rho) for the association between SD_BOLD_ and SD_EEG_ for A) younger (N=135) and B) older (N=54) adults for different frequency bands across each pair of 96 regions of interests. The correlations between SD_BOLD_ and SD_EEG_ were tested using pairwise Spearman’s rank correlation corrected for multiple comparison by false discovery rates (FDR; Benjamini and Hochberg, 1995).

**Figure 5.**
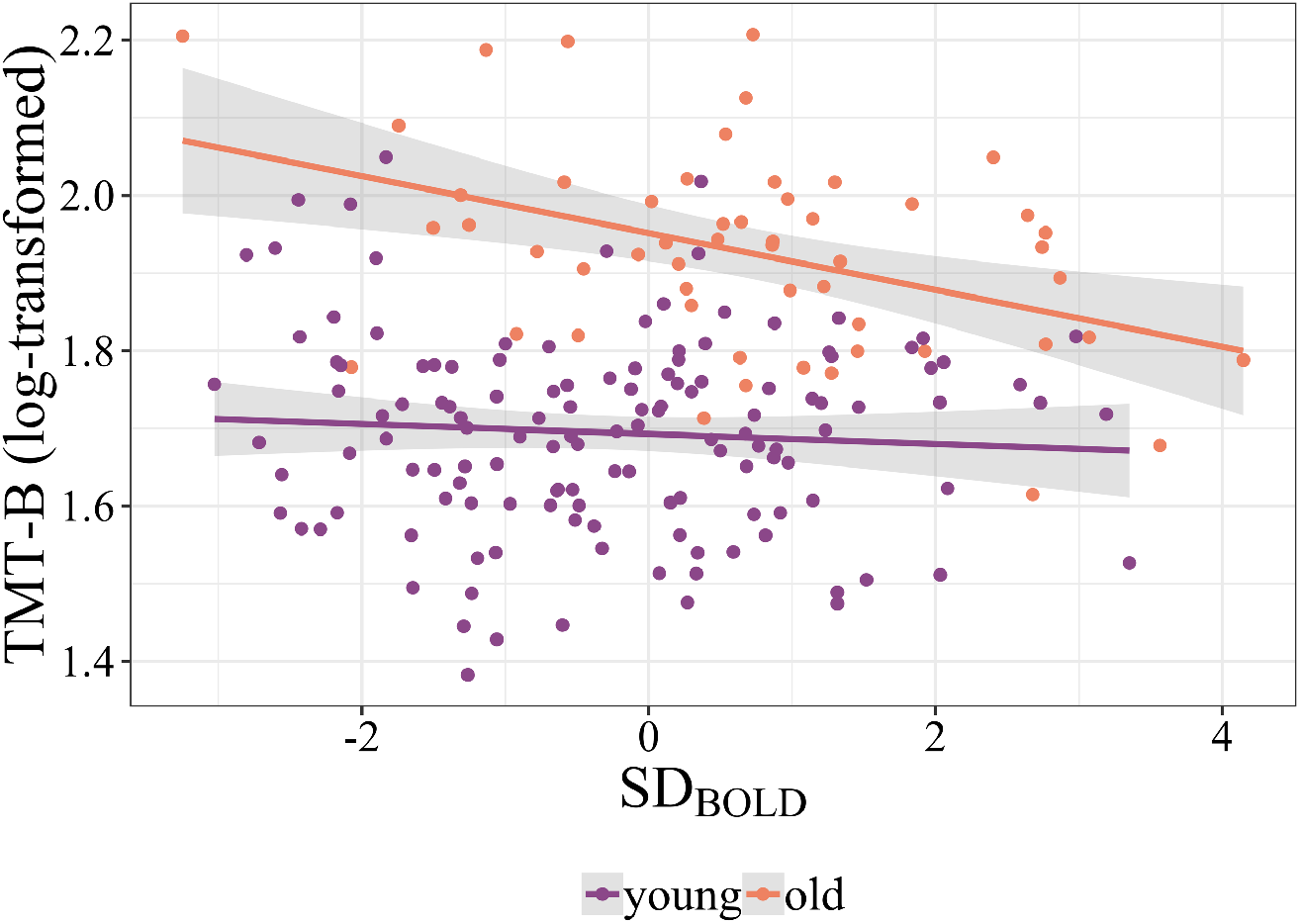
Age-dependent relationship between cognitive performance and BOLD signal variability. The scatterplot shows the significant association between task completion time in TMT-B (x-axis) and SD_BOLD_ (adjusted R^2^ = 0.395, *F*(7,181) = 18.60, p <.001, interaction: β= −0.002, p = 0.027) in PC2, driven mainly by the left anterior and posterior temporal gyrus, bilateral anterior and posterior cingulate cortex.

## 4. Discussion

Comparing healthy younger and older adults, we found widespread variability reductions in BOLD signal as well as in the amplitude envelope of delta, theta, and alpha frequency of rsEEG, whereas increased variability with aging was observed in the beta-band frequency. As a complementary analysis, we also explored sex differences and found that female subjects exhibited higher EEG signal variability than male subjects; no significant sex differences were found in BOLD signal variability. There were no significant correlations between hemodynamic (SD_BOLD_) and electrophysiological (SD_EEG_) measures of brain signal variability, neither in the younger nor in the older adults. Our results suggest that variability measures of rsfMRI and rsEEG – while both related to aging – are dominated by different physiological origins and relate differently to age and sex.

### 4.1. BOLD Signal Variability

The first aim of our study was to investigate the effect of age on BOLD signal variability, as measured by SD of spontaneous fluctuations during rsfMRI. Consistent with recent rsfMRI studies demonstrating that BOLD signal variability decreases with age in large-scale networks (Grady and Garrett, 2018; Nomi et al., 2017), we found that older subjects had reduced SD_BOLD_ in temporal and occipital brain regions but also in cortical midline structures like the precuneus, anterior and posterior cingulate cortices, as well as orbitofrontal cortex compared to younger adults. These age-related reductions in BOLD signal variability were thus especially apparent in regions of the Default Mode (DMN) and the Fronto-Parietal Network (FPN). The DMN is an intrinsically correlated network of brain regions, that is particularly active during rest or fixation blocks (Biswal et al., 2010). It reflects the systematic integration of information across the cortex (Margulies et al., 2016) and has been frequently associated with psychological functions like self-referential thought or mind-wandering, and also memory retrieval (Andrews-Hanna et al., 2014; Raichle, 2015). The FPN is involved in cognitive control processes (Spreng et al., 2013), and closely interacts with the DMN, for example during mind-wandering state (Golchert et al., 2017). Previous studies in healthy aging noted that older subjects showed lower functional connectivity in DMN and FPN regions (Damoiseaux, 2017; Damoiseaux et al., 2008; Meunier et al., 2009; Petersen et al., 2014). Similarly, an altered functional connectivity in the DMN has been found in different pathologies, for example, in Alzheimer’s disease (Greicius et al., 2004) or mild cognitive impairment (Das et al., 2015). Further, we found a significant interaction between age and SD_BOLD_ in temporal and cingulate cortices for performance on the cognitive task (TMT-B), suggesting that the relationship between brain signal variability and cognitive performance depends on the participants’ age. We speculate that – in the elderly – reduced BOLD signal variability in the DMN and the FPN, particularly in the overlapping frontal brain regions, could be related to locally impaired function that is reflected in impaired cognitive performance (Campbell et al., 2012). Such findings support the notion that local BOLD signal variability may be a valuable biomarker of neurocognitive health (and disease) in aging.

Sex-specific differences in brain structure and function have been previously shown (for a review see, Gong et al., 2011; Ruigrok et al., 2014; Sacher et al., 2013). For example, larger total brain volume has been reported in male as compared to female subjects (Gong et al., 2011), whereas higher cerebral blood flow (Gur et al., 1982; Rodriguez et al., 1988) and stronger functional connectivity in the DMN (Tomasi and Volkow, 2012) were found in females. In our exploratory analysis, we did not find significant sex differences in BOLD signal variability when controlling for total intracranial volume as an approximation of overall brain size.

### 4.2. Electrophysiological Signal Variability

Measures of neural variability were derived from rsEEG for several main frequency bands (delta, theta, alpha, beta) as the standard deviation of their amplitude of envelope time series data, analogously to the BOLD signal variability. Multimodal imaging studies have shown that the amplitude envelope of neural oscillatory activity across frequency bands relates to different rsfMRI networks (Brookes et al., 2011; Deligianni et al., 2014), confirming the neurophysiological origin of the resting state networks measured with BOLD fMRI. Additionally, these studies also concluded that different frequency bands can be related to the same functional network, but also differentially to distinct networks (Brookes et al., 2011; Laufs et al., 2006; Mantini et al., 2007; Meyer et al., 2013). For instance, Mantini et. al. (2007) reported that the visual network is associated with all frequency bands except gamma rhythm, while the sensorimotor network is primarily associated with beta-band oscillations.

In our analysis, we found age-dependent EEG signal variability changes within networks which were associated with more than one frequency band, thus confirming that neurons generating oscillations at different frequencies may contribute to the same network. More precisely, we found age-related reductions in SD_DELTA_ and SD_ALPHA_ mainly in a visual network (including calcarine regions, cuneal cortex, and occipital pole), SD_THETA_ in posterior DMN (e.g., posterior cingulate cortex), while an enhancement of SD_BETA_ was mainly seen in the temporal (e.g., superior/middle temporal gyrus), and central/sensorimotor (e.g., supramarginal gyrus) regions. These results align with previous reports of age-dependent changes of electrophysiological activity using spectral power (Dustman et al., 1993; Vlahou et al., 2015), and signal variability (Dustman et al., 1999; Tsvetanov et al., 2015).

Age-related decreases of alpha amplitude and alpha band variability (measured by SD of the oscillatory signal) were previously found in posterior and occipital brain regions (Babiloni et al., 2006; Tsvetanov et al., 2015). Alpha rhythm is a classical EEG hallmark of resting wakefulness (Laufs et al., 2003) that is modulated by thalamo-cortical and cortico-cortical interactions (Bazanova and Vernon, 2014; Goldman et al., 2002; Lopes Da Silva et al., 1997; Moosmann et al., 2003). It has been suggested that the posterior alpha-frequency plays an important role in the top-down control of cortical activation and excitability (Klimesch, 1999). Accordingly, decreased alpha variability in occipital regions might be associated with altered functioning of the cholinergic basal forebrain, affecting thalamo-cortical and cortico-cortical processing. Our finding of higher temporal and sensorimotor SD_BETA_ in the elderly is in line with previous findings (Rossiter et al., 2014; Tsvetanov et al., 2015). Aging has previously been associated with an increase in movement-related beta-band attenuation, suggesting an enhanced motor cortex GABAergic inhibitory activity in older individuals (Rossiter et al., 2014). Similarly, beta-band activity is thought to play a key role in signaling maintenance of the status quo of the motor system, despite the absence of movement (Engel and Fries, 2010). Therefore, greater SD_BETA_ in sensorimotor brain regions could be interpreted as a compensatory mechanism to account for a decline of motor performance during aging (Quandt et al., 2016).

It should be noted that the present findings of age-related alterations of brain signal variability at different frequencies might be influenced by several anatomical factors which might influence EEG-generators such as reduced cortical gray matter (Babiloni et al., 2013; Moretti et al., 2012), white-matter (Nunez et al., 2015; Valdés-Hernández et al., 2010), and increased amount of cerebrospinal fluid (CSF; Hartikainen et al., 1992; Stomrud et al., 2010), but also alterations of cerebral glucose metabolism (Dierks et al., 2000). Localized or global disturbances of brain anatomy and function might lead to deviations in the EEG sources, resulting in EEG amplitude changes. A methodological improvement for future studies will therefore be the application of individual head models (Ziegler et al., 2014).

In addition to the effect of age on rsEEG signal variability, an exploratory analysis showed sex differences in distinct brain regions and EEG frequencies. More precisely, we found higher SD_DELTA_ and SD_THETA_ in occipito-temporal, SD_ALPHA_ in frontal, and SD_BETA_ in frontal as well as occipito-temporal brain regions in female compared to male subjects. While some studies demonstrated higher alpha (Aurlien et al., 2003), delta (Armitage, 1995), theta (Carrier et al., 2001; Duffy et al., 1993), and beta power (Jaušovec and Jaušovec, 2010; Matsuura et al., 1985; Veldhuizen et al., 1993) in female relative to male subjects, other studies reported the opposite pattern (Brenner et al., 1995; Latta et al., 2005; Zappasodi et al., 2006). These differences in EEG signal variability could be a result of different mechanisms (biological/hormonal, cultural or developmental) involved in shaping sex differences. Unfortunately, based on our dataset we cannot differentiate which of these potential mechanisms might be most relevant for the observed changes.

### 4.3. Association between BOLD and EEG Variability

We further assessed how neural variability in source-reconstructed rsEEG related to the analogous parameters of BOLD signal variability in rsfMRI using univariate and multivariate correlation analyses. Previously, simultaneous EEG-fMRI studies have shown meaningful relationships between fluctuations in EEG power, frequency, phase, and local BOLD changes (for a review see, Jorge et al., 2014; Ritter and Villringer, 2006). Due to age-related physiological (particularly cardiovascular) alterations in the brain, we expected the relationship between BOLD and EEG signal variability to be stronger in younger than older adults. However, in the present study, both univariate and multivariate analyses showed no significant correlations between SD_BOLD_ and SD_EEG_ neither in the younger nor in the older adults. This finding was supported by the distinct anatomical distributions of age-related changes in BOLD and EEG signal variability, that barely showed a spatial overlap, suggesting different underlying physiological processes. What could they be? Clearly, neuronal activity is the main signal source for EEG- and MEG recordings as well as for EEG/MEG-based variability measures. BOLD signal variability, however, can reflect both vascular and neural processes (Garrett et al., 2017). While neuronal activity clearly contributes to the BOLD signal at rest (Ma et al., 2016; Mateo et al., 2017), our results indicate, however, that neuronal activity which is captured by EEG (or more specifically by our EEG-based measures), is not the major determinant of BOLD variability in the resting state. Other factors that could contribute to BOLD variability are (i) neuronal activity which is not captured by EEG and (ii) non-neural factors such as vasomotion, or cardiac and respiratory signals (Murphy et al., 2013). In the elderly, additional factors related to the known morphological and functional changes of blood vessels as well as age-related metabolic changes are known to affect CBF (Ances et al., 2009; Martin et al., 1991), CMRO_2_ (Aanerud et al., 2012), and CVR (Liu et al., 2013) and therefore are likely to also influence BOLD variability. Thus, given different underlying physiology, joint EEG and fMRI variability studies might provide complementary information for a comprehensive assessment of neuronal as well as vascular factors related to aging.

## 5. Limitations

There are several limitations of our study: EEG and MRI scans were not recorded simultaneously. Therefore, we could not directly relate the two signals in a cross-correlation analysis. Furthermore, EEG and MRI were performed with different body postures (fMRI; supine, EEG; seated) known to affect brain function, for example, changes in the amplitude of the EEG signal have been related to different body postures presumably due to the shifts in cerebrospinal fluid layer thickness (Rice et al., 2013). Similarly, other experimental (e.g., visual display; Nir et al., 2006), environmental (e.g., acoustic noise in MRI; Andoh et al., 2017; Cho et al., 1998) and subject-related factors (e.g., changes of vigilance; Tagliazucchi and Laufs, 2014; Wong et al., 2013) could have introduced unintended variations in our results (Yan et al., 2013) and the influence of these factors is probably not the same for the different methods, e.g., noise in MRI or poor “control” of vigilance in MRI. For instance, given the well-known relationship between vigilance or arousal and fMRI signal fluctuations (Bijsterbosch et al., 2017; Chang et al., 2016; Haimovici et al., 2017), it is likely that the observed age-related differences in BOLD signal variability might be confounded by such within-subject (state) variability. Therefore, future rsfMRI studies may benefit from obtaining arousal-related (e.g., self-report) measures and an explicit measurement of eye movements and eye opening/closure to account for the influence of arousal on the BOLD amplitude changes. Another option would be to combine EEG and fMRI simultaneously. Yet, resting state measures of EEG (Näpflin et al., 2007) and fMRI (Shehzad et al., 2009; Zuo et al., 2010) have been shown to be reliable within-individuals across time.

In our study, the computation of the source reconstructed rsEEG required the parcellation of the brain into relatively large anatomical ROIs. It could well be that the analysis with a higher spatial resolution (e.g., at the voxel-level) with individual head models may provide additional insights about brain signal variability.

Finally, while our study aimed at comparing analogous variability measures in EEG and fMRI, future research using rsEEG and rsfMRI in the same subjects would benefit from the addition of connectivity-based measures including graph theory-based (Yu et al., 2016) or sliding-window methods (Chang et al., 2013; Qin et al., 2019).

## 6. Conclusion

In this study, we report age and sex differences of brain signal variability obtained with rsfMRI and rsEEG from the same subjects. We demonstrate extensive age-related reduction of SD_BOLD_, SD_DELTA_, SD_THETA_, and SD_ALPHA_ mainly in the DMN and the visual network, while a significant increase of SD_BETA_ was mainly seen in temporal brain regions. We could not demonstrate significant associations between SD_BOLD_ and SD_EEG._ Our findings indicate that measurements of BOLD and EEG signal variability, respectively, are likely to stem from different physiological origins and relate differentially to age and sex. While the two types of measurements are thus not interchangeable, it seems, however, plausible that both markers of brain variability may provide complementary information about the aging process.

## 8. Acknowledgements

We gratefully acknowledge the Mind-Body- Emotion group at the Max Planck Institute for Human Cognitive and Brain Sciences.

