## Supplementary Figure 1 for "BOLD and EEG Signal Variability at Rest Differently Relate to Aging in the Human Brain"

### 2. Supplementary Material

**Supplementary Figure 1.** Flowchart of selecting participants from the Mind-Brain-Body study.

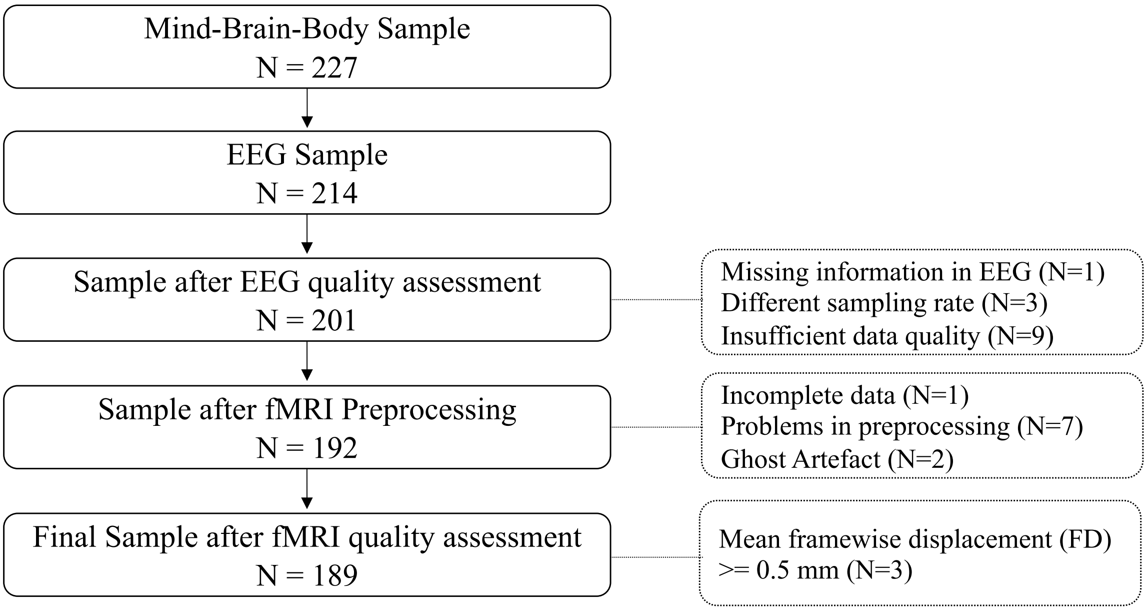

**Supplementary Figure 2.** Spatial maps of mean $SD_{BOLD}$ and $SD_{EEG}$ for younger adults (N=135). We computed the mean $SD_{BOLD}$ and $SD_{EEG}$ by I) log-transforming the SD values, II) averaging separately for younger and older subjects, and III) then back-transforming the values (McDonald, 2014).

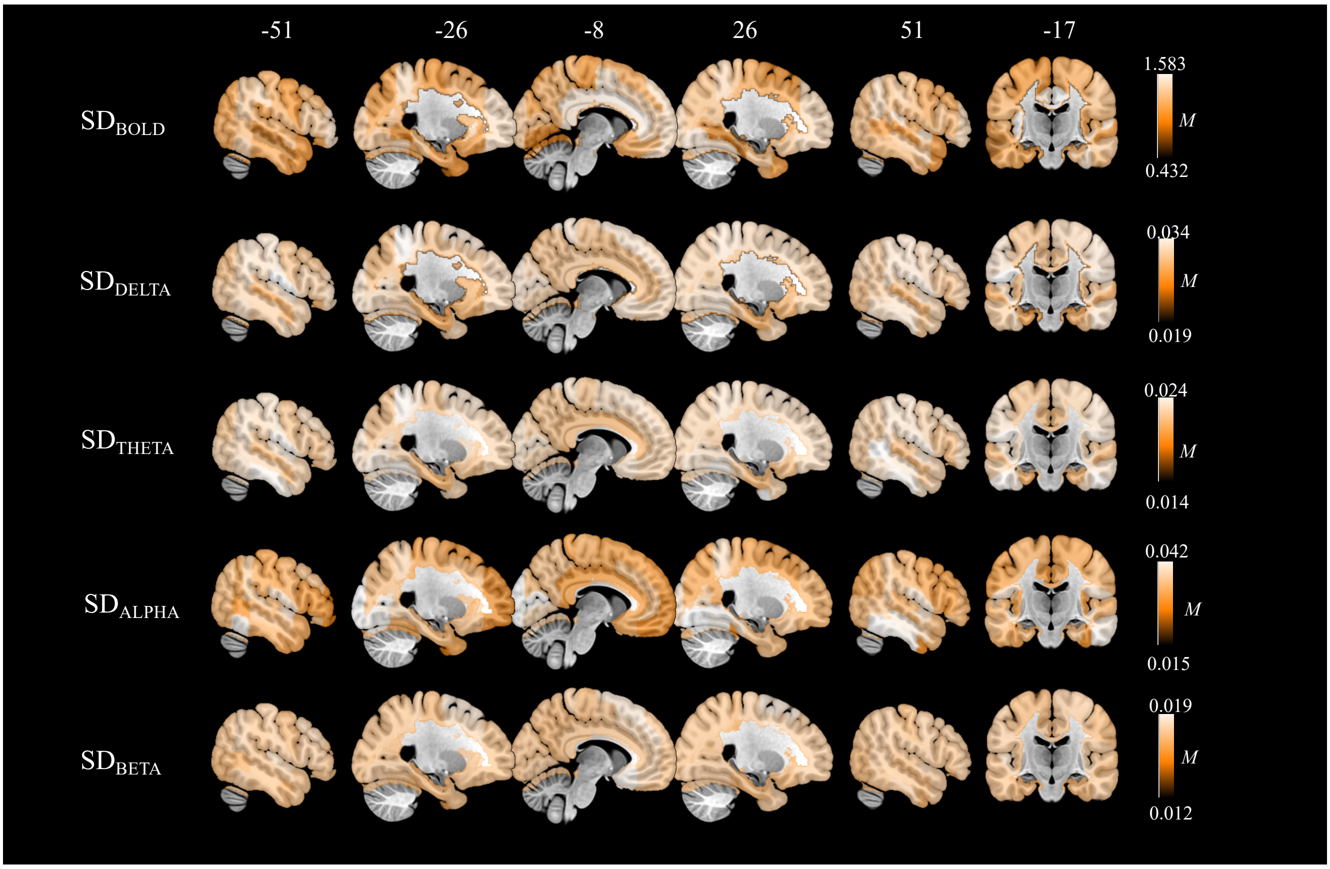

**Supplementary Figure 3.** Spatial maps of mean $SD_{BOLD}$ and $SD_{EEG}$ for older adults (N=54). We computed the mean $SD_{BOLD}$ and $SD_{EEG}$ by I) log-transforming the SD values, II) averaging separately for younger and older subjects, and III) then back-transforming the values (McDonald, 2014).

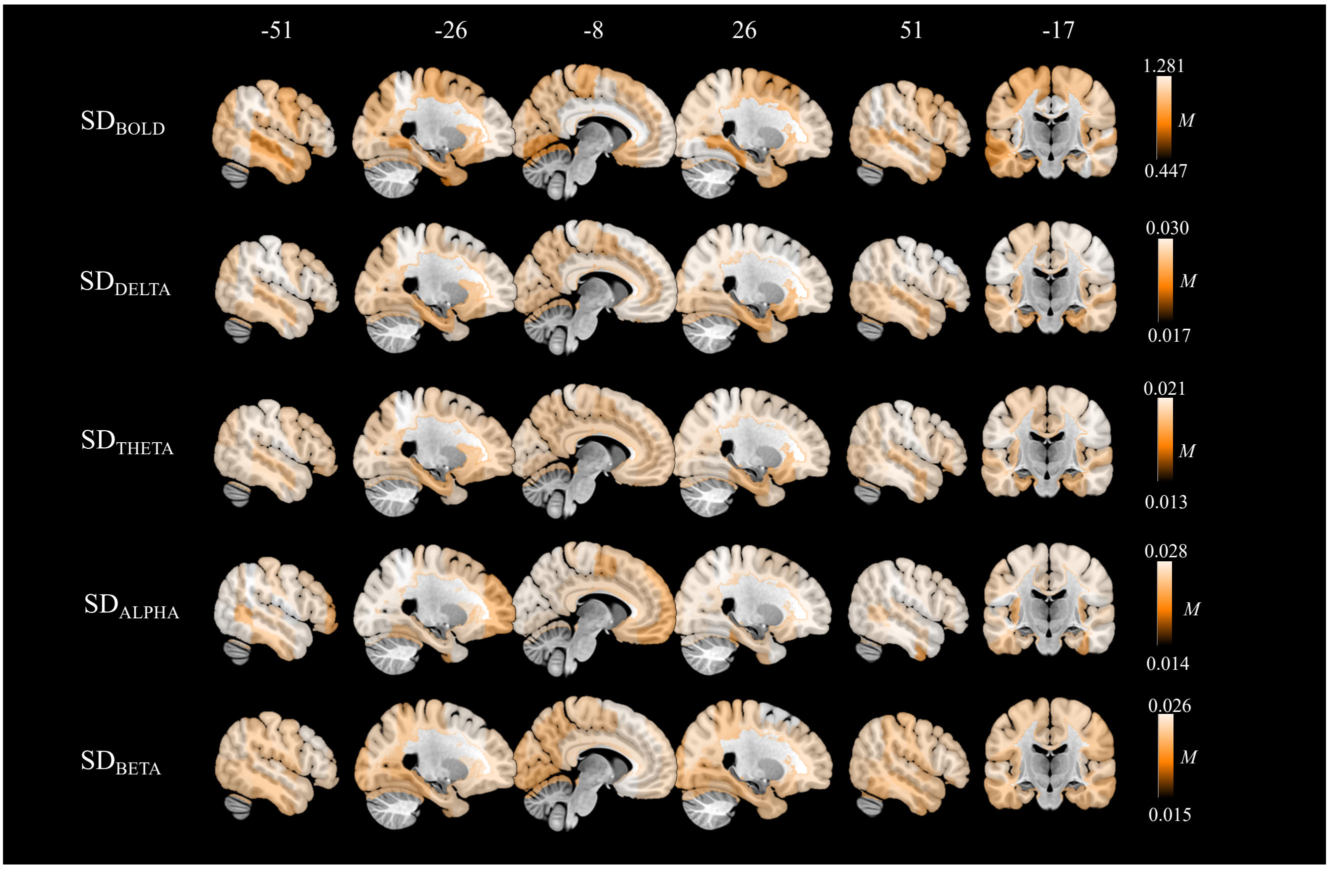

**Supplementary Figure 4.** Graphical distribution of the F-values for significant effects of age group or sex. The nonparametric ANCOVAs with $SD_{BOLD}$ as dependent variable demonstrated that there was a significant main effect of age group in 72 ROIs, while no significant effect of sex was observed. We further found significant main effects of age group in all EEG frequency bands: $SD_{DELTA}$ in 14 ROIs (F-values: 12.57–20.94), $SD_{THETA}$ in 16 ROIs (F-values: 13.16–40.30), $SD_{ALPHA}$ in 20 ROIs (F-values: 12.69–20.12), and $SD_{BETA}$ in 69 ROIs (F-values: 12.50–21.61). There were also significant main effects of sex in all frequency bands: $SD_{DELTA}$ in 21 ROIs (F-values: 13.24–26.63), $SD_{THETA}$ in 74 ROIs (F-values: 12.68–30.06), $SD_{ALPHA}$ in 4 ROIs (F-values: 12.88–16.51), and $SD_{BETA}$ in 69 ROIs (F-values: 12.54–35.72).

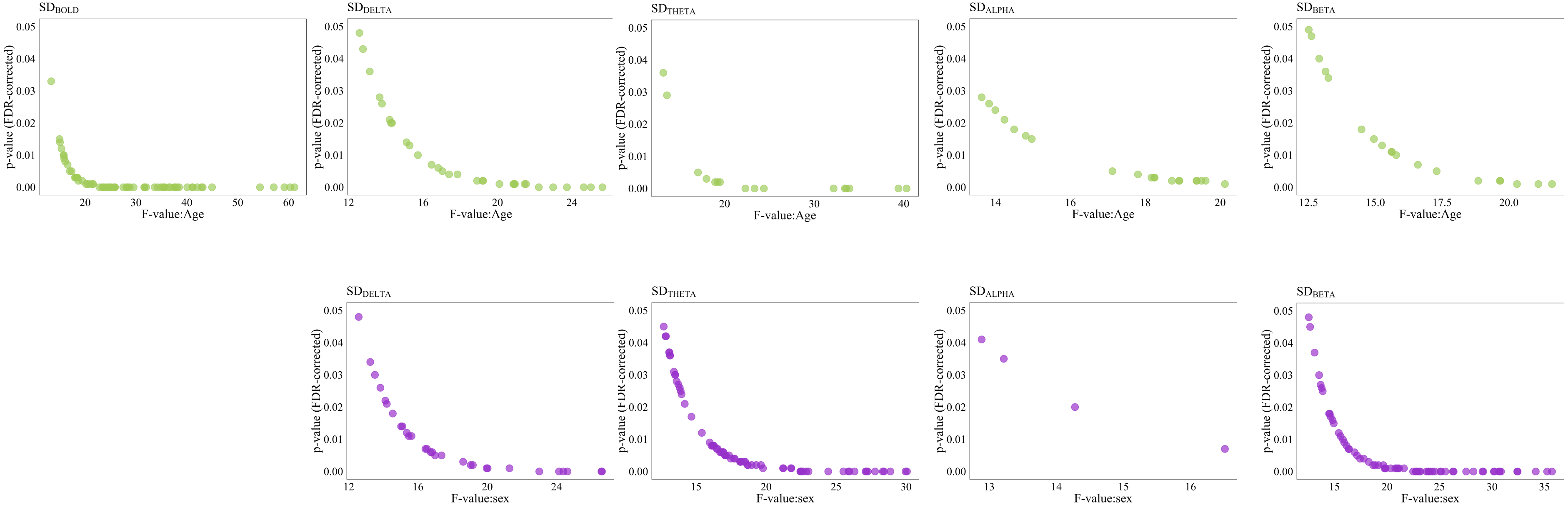

**Supplementary Table 1:** Table showing the mean (*M*) values of the $SD_{BOLD}$ and $SD_{EEG}$ for younger (N=135) and older (N=54) adults, respectively.

| | $SD_{BOLD}$ | | $SD_{DELTA}$ | | $SD_{THETA}$ | | $SD_{ALPHA}$ | | $SD_{BETA}$ | |
|---|---|---|---|---|---|---|---|---|---|---|
| **ROI** | **Young** | **Old** | **Young** | **Old** | **Young** | **Old** | **Young** | **Old** | **Young** | **Old** |
| Left Frontal Pole | 1.336 | 1.085 | 0.028 | 0.027 | 0.019 | 0.016 | 0.015 | 0.016 | 0.014 | 0.021 |
| Right Frontal Pole | 1.304 | 1.029 | 0.027 | 0.027 | 0.020 | 0.018 | 0.026 | 0.026 | 0.015 | 0.021 |
| Left Insular Cortex | 1.583 | 1.213 | 0.023 | 0.019 | 0.021 | 0.014 | 0.022 | 0.016 | 0.014 | 0.018 |
| Right Insular Cortex | 0.845 | 0.728 | 0.024 | 0.022 | 0.019 | 0.015 | 0.021 | 0.018 | 0.016 | 0.023 |
| Left Superior Frontal Gyrus | 0.883 | 0.767 | 0.030 | 0.029 | 0.021 | 0.018 | 0.021 | 0.021 | 0.018 | 0.024 |
| Right Superior Frontal Gyrus | 0.735 | 0.651 | 0.028 | 0.029 | 0.021 | 0.018 | 0.022 | 0.023 | 0.018 | 0.026 |
| Left Middle Frontal Gyrus | 0.981 | 0.848 | 0.024 | 0.023 | 0.020 | 0.016 | 0.029 | 0.022 | 0.016 | 0.024 |
| Right Middle Frontal Gyrus | 0.868 | 0.789 | 0.029 | 0.030 | 0.020 | 0.019 | 0.021 | 0.024 | 0.015 | 0.020 |
| Left Inferior Frontal Gyrus, pars triangularis | 1.170 | 0.992 | 0.026 | 0.027 | 0.017 | 0.017 | 0.019 | 0.024 | 0.015 | 0.023 |
| Right Inferior Frontal Gyrus, pars triangularis | 0.977 | 0.848 | 0.024 | 0.025 | 0.016 | 0.016 | 0.020 | 0.024 | 0.014 | 0.022 |
| Left Inferior Frontal Gyrus, pars opercularis | 1.162 | 0.905 | 0.027 | 0.028 | 0.018 | 0.017 | 0.020 | 0.024 | 0.015 | 0.022 |
| Right Inferior Frontal Gyrus, pars opercularis | 1.077 | 0.949 | 0.025 | 0.026 | 0.017 | 0.017 | 0.021 | 0.025 | 0.014 | 0.022 |
| Left Precentral Gyrus | 0.710 | 0.646 | 0.024 | 0.023 | 0.017 | 0.016 | 0.024 | 0.025 | 0.013 | 0.019 |
| Right Precentral Gyrus | 0.875 | 0.766 | 0.028 | 0.028 | 0.019 | 0.018 | 0.022 | 0.025 | 0.015 | 0.020 |
| Left Temporal Pole | 0.733 | 0.741 | 0.027 | 0.026 | 0.019 | 0.018 | 0.023 | 0.026 | 0.014 | 0.021 |
| Right Temporal Pole | 0.761 | 0.746 | 0.027 | 0.025 | 0.019 | 0.018 | 0.027 | 0.027 | 0.015 | 0.019 |
| Left Superior Temporal Gyrus, anterior division | 1.089 | 1.028 | 0.022 | 0.021 | 0.017 | 0.015 | 0.029 | 0.023 | 0.015 | 0.023 |
| Right Superior Temporal Gyrus, anterior division | 1.041 | 0.921 | 0.020 | 0.017 | 0.016 | 0.013 | 0.030 | 0.022 | 0.013 | 0.020 |
| Left Superior Temporal Gyrus, posterior division | 0.485 | 0.447 | 0.019 | 0.019 | 0.015 | 0.014 | 0.025 | 0.022 | 0.014 | 0.022 |
| Right Superior Temporal Gyrus, posterior division | 0.748 | 0.693 | 0.020 | 0.019 | 0.015 | 0.014 | 0.026 | 0.023 | 0.013 | 0.020 |
| Left Middle Temporal Gyrus, anterior division | 0.457 | 0.507 | 0.020 | 0.019 | 0.015 | 0.014 | 0.026 | 0.023 | 0.012 | 0.019 |
| Right Middle Temporal Gyrus, anterior division | 1.255 | 1.019 | 0.024 | 0.019 | 0.017 | 0.014 | 0.033 | 0.023 | 0.013 | 0.017 |
| Left Middle Temporal Gyrus, posterior division | 0.432 | 0.458 | 0.026 | 0.022 | 0.019 | 0.016 | 0.029 | 0.024 | 0.014 | 0.017 |
| Right Middle Temporal Gyrus, posterior division | 1.148 | 1.078 | 0.029 | 0.024 | 0.021 | 0.019 | 0.038 | 0.026 | 0.014 | 0.016 |

| | | | | | | | | | | |
|---|---|---|---|---|---|---|---|---|---|---|
| Left Middle Temporal Gyrus, temporooccipital part | 1.009 | 0.888 | 0.027 | 0.028 | 0.020 | 0.018 | 0.017 | 0.016 | 0.012 | 0.018 |
| Right Middle Temporal Gyrus, temporooccipital part | 0.776 | 0.720 | 0.026 | 0.022 | 0.024 | 0.017 | 0.029 | 0.020 | 0.015 | 0.019 |
| Left Inferior Temporal Gyrus, anterior division | 0.819 | 0.773 | 0.028 | 0.028 | 0.022 | 0.019 | 0.022 | 0.020 | 0.013 | 0.017 |
| Right Inferior Temporal Gyrus, anterior division | 0.982 | 0.821 | 0.024 | 0.022 | 0.020 | 0.015 | 0.017 | 0.014 | 0.012 | 0.016 |
| Left Inferior Temporal Gyrus, posterior division | 0.886 | 0.737 | 0.026 | 0.023 | 0.023 | 0.017 | 0.025 | 0.019 | 0.014 | 0.017 |
| Right Inferior Temporal Gyrus, posterior division | 1.052 | 0.891 | 0.030 | 0.024 | 0.023 | 0.019 | 0.040 | 0.026 | 0.015 | 0.019 |
| Left Inferior Temporal Gyrus, temporooccipital part | 1.224 | 1.080 | 0.029 | 0.022 | 0.021 | 0.018 | 0.040 | 0.026 | 0.015 | 0.019 |
| Right Inferior Temporal Gyrus, temporooccipital part | 1.052 | 0.947 | 0.029 | 0.022 | 0.021 | 0.018 | 0.041 | 0.025 | 0.014 | 0.016 |
| Left Postcentral Gyrus | 0.940 | 0.845 | 0.030 | 0.029 | 0.021 | 0.019 | 0.021 | 0.022 | 0.015 | 0.020 |
| Right Postcentral Gyrus | 1.089 | 0.946 | 0.030 | 0.029 | 0.021 | 0.020 | 0.026 | 0.025 | 0.013 | 0.017 |
| Left Superior Parietal Lobule | 1.309 | 1.174 | 0.033 | 0.029 | 0.023 | 0.021 | 0.033 | 0.028 | 0.014 | 0.017 |
| Right Superior Parietal Lobule | 1.282 | 1.093 | 0.030 | 0.026 | 0.021 | 0.020 | 0.035 | 0.027 | 0.014 | 0.017 |
| Left Supramarginal Gyrus, anterior division | 1.291 | 1.127 | 0.029 | 0.029 | 0.020 | 0.019 | 0.023 | 0.025 | 0.014 | 0.018 |
| Right Supramarginal Gyrus, anterior division | 1.213 | 1.051 | 0.030 | 0.027 | 0.021 | 0.019 | 0.027 | 0.027 | 0.014 | 0.019 |
| Left Supramarginal Gyrus, posterior division | 1.148 | 1.092 | 0.030 | 0.027 | 0.021 | 0.019 | 0.032 | 0.028 | 0.015 | 0.018 |
| Right Supramarginal Gyrus, posterior division | 1.100 | 0.968 | 0.029 | 0.025 | 0.021 | 0.019 | 0.032 | 0.026 | 0.015 | 0.017 |
| Left Angular Gyrus | 1.230 | 1.076 | 0.028 | 0.028 | 0.021 | 0.018 | 0.023 | 0.023 | 0.016 | 0.022 |
| Right Angular Gyrus | 1.323 | 1.149 | 0.025 | 0.026 | 0.018 | 0.017 | 0.024 | 0.025 | 0.015 | 0.022 |
| Left Lateral Occipital Cortex, superior division | 0.811 | 0.765 | 0.023 | 0.022 | 0.017 | 0.016 | 0.028 | 0.026 | 0.014 | 0.021 |
| Right Lateral Occipital Cortex, superior division | 1.072 | 0.981 | 0.027 | 0.028 | 0.019 | 0.018 | 0.022 | 0.025 | 0.014 | 0.021 |
| Left Lateral Occipital Cortex, inferior division | 0.890 | 0.861 | 0.026 | 0.025 | 0.019 | 0.018 | 0.027 | 0.027 | 0.013 | 0.019 |
| Right Lateral Occipital Cortex, inferior division | 1.100 | 0.892 | 0.024 | 0.023 | 0.018 | 0.016 | 0.027 | 0.026 | 0.013 | 0.020 |
| Left Intracalcarine Cortex | 1.072 | 0.863 | 0.030 | 0.023 | 0.021 | 0.018 | 0.041 | 0.026 | 0.014 | 0.016 |
| Right Intracalcarine Cortex | 0.885 | 0.893 | 0.028 | 0.023 | 0.020 | 0.017 | 0.034 | 0.024 | 0.014 | 0.015 |
| Left Frontal Medial Cortex | 0.946 | 0.945 | 0.029 | 0.027 | 0.019 | 0.016 | 0.017 | 0.016 | 0.016 | 0.022 |
| Right Frontal Medial Cortex | 1.349 | 1.063 | 0.027 | 0.026 | 0.020 | 0.017 | 0.028 | 0.024 | 0.015 | 0.021 |
| Left Juxtapositional Lobule Cortex (formerly Supplementary Motor Cortex) | 1.380 | 1.102 | 0.023 | 0.019 | 0.020 | 0.014 | 0.022 | 0.016 | 0.014 | 0.018 |

| | | | | | | | | | | |
|---|---|---|---|---|---|---|---|---|---|---|
| Right Juxtapositional Lobule Cortex (formerly Supplementary Motor Cortex) | 0.925 | 0.784 | 0.024 | 0.020 | 0.019 | 0.014 | 0.023 | 0.018 | 0.017 | 0.023 |
| Left Subcallosal Cortex | 0.880 | 0.762 | 0.030 | 0.028 | 0.021 | 0.017 | 0.023 | 0.020 | 0.019 | 0.025 |
| Right Subcallosal Cortex | 0.984 | 0.846 | 0.028 | 0.027 | 0.021 | 0.017 | 0.024 | 0.021 | 0.019 | 0.025 |
| Left Paracingulate Gyrus | 1.251 | 0.902 | 0.023 | 0.021 | 0.019 | 0.015 | 0.029 | 0.022 | 0.016 | 0.023 |
| Right Paracingulate Gyrus | 0.942 | 0.822 | 0.029 | 0.028 | 0.019 | 0.018 | 0.021 | 0.022 | 0.015 | 0.021 |
| Left Cingulate Gyrus, anterior division | 1.487 | 1.213 | 0.024 | 0.026 | 0.015 | 0.016 | 0.020 | 0.022 | 0.014 | 0.022 |
| Right Cingulate Gyrus, anterior division | 1.459 | 1.201 | 0.023 | 0.024 | 0.015 | 0.016 | 0.022 | 0.024 | 0.013 | 0.020 |
| Left Cingulate Gyrus, posterior division | 1.412 | 1.163 | 0.025 | 0.027 | 0.016 | 0.016 | 0.020 | 0.023 | 0.014 | 0.021 |
| Right Cingulate Gyrus, posterior division | 1.579 | 1.281 | 0.024 | 0.025 | 0.016 | 0.016 | 0.023 | 0.025 | 0.014 | 0.020 |
| Left Precuneous Cortex | 0.978 | 0.878 | 0.024 | 0.021 | 0.017 | 0.015 | 0.029 | 0.025 | 0.013 | 0.016 |
| Right Precuneous Cortex | 1.120 | 1.041 | 0.027 | 0.027 | 0.018 | 0.017 | 0.022 | 0.023 | 0.014 | 0.019 |
| Left Cuneal Cortex | 0.793 | 0.764 | 0.027 | 0.026 | 0.018 | 0.017 | 0.027 | 0.026 | 0.014 | 0.019 |
| Right Cuneal Cortex | 0.894 | 0.837 | 0.028 | 0.024 | 0.020 | 0.017 | 0.033 | 0.027 | 0.015 | 0.017 |
| Left Frontal Orbital Cortex | 0.813 | 0.702 | 0.021 | 0.020 | 0.017 | 0.014 | 0.029 | 0.023 | 0.014 | 0.021 |
| Right Frontal Orbital Cortex | 1.166 | 0.850 | 0.021 | 0.017 | 0.016 | 0.013 | 0.030 | 0.022 | 0.013 | 0.018 |
| Left Parahippocampal Gyrus, anterior division | 0.853 | 0.733 | 0.019 | 0.018 | 0.014 | 0.013 | 0.027 | 0.023 | 0.013 | 0.020 |
| Right Parahippocampal Gyrus, anterior division | 1.129 | 0.975 | 0.020 | 0.017 | 0.015 | 0.013 | 0.029 | 0.023 | 0.013 | 0.018 |
| Left Parahippocampal Gyrus, posterior division | 1.101 | 0.943 | 0.021 | 0.017 | 0.016 | 0.013 | 0.030 | 0.022 | 0.013 | 0.016 |
| Right Parahippocampal Gyrus, posterior division | 0.459 | 0.455 | 0.024 | 0.018 | 0.018 | 0.014 | 0.035 | 0.022 | 0.013 | 0.016 |
| Left Lingual Gyrus | 0.550 | 0.554 | 0.027 | 0.021 | 0.019 | 0.016 | 0.034 | 0.024 | 0.015 | 0.015 |
| Right Lingual Gyrus | 0.558 | 0.544 | 0.030 | 0.023 | 0.021 | 0.019 | 0.041 | 0.026 | 0.015 | 0.016 |
| Left Temporal Fusiform Cortex, anterior division | 0.524 | 0.507 | 0.027 | 0.028 | 0.020 | 0.018 | 0.017 | 0.017 | 0.012 | 0.018 |
| Right Temporal Fusiform Cortex, anterior division | 0.929 | 0.807 | 0.026 | 0.021 | 0.024 | 0.016 | 0.029 | 0.020 | 0.015 | 0.019 |
| Left Temporal Fusiform Cortex, posterior division | 0.870 | 0.763 | 0.028 | 0.027 | 0.022 | 0.018 | 0.022 | 0.020 | 0.013 | 0.017 |
| Right Temporal Fusiform Cortex, posterior division | 1.387 | 1.244 | 0.024 | 0.022 | 0.020 | 0.015 | 0.017 | 0.015 | 0.013 | 0.017 |
| Left Temporal Occipital Fusiform Cortex | 1.258 | 0.961 | 0.027 | 0.023 | 0.023 | 0.017 | 0.025 | 0.019 | 0.014 | 0.018 |
| Right Temporal Occipital Fusiform Cortex | 1.367 | 1.187 | 0.031 | 0.024 | 0.023 | 0.019 | 0.040 | 0.027 | 0.015 | 0.019 |

| | | | | | | | | | | |
|---|---|---|---|---|---|---|---|---|---|---|
| Left Occipital Fusiform Gyrus | 1.182 | 0.974 | 0.029 | 0.022 | 0.022 | 0.018 | 0.041 | 0.026 | 0.015 | 0.018 |
| Right Occipital Fusiform Gyrus | 1.416 | 1.169 | 0.029 | 0.022 | 0.021 | 0.018 | 0.042 | 0.025 | 0.015 | 0.016 |
| Left Frontal Operculum Cortex | 1.039 | 0.953 | 0.030 | 0.029 | 0.021 | 0.019 | 0.022 | 0.021 | 0.016 | 0.022 |
| Right Frontal Operculum Cortex | 1.299 | 1.207 | 0.031 | 0.028 | 0.021 | 0.019 | 0.027 | 0.025 | 0.014 | 0.018 |
| Left Central Opercular Cortex | 1.012 | 0.930 | 0.034 | 0.029 | 0.023 | 0.020 | 0.036 | 0.028 | 0.015 | 0.017 |
| Right Central Opercular Cortex | 1.032 | 0.976 | 0.031 | 0.026 | 0.022 | 0.020 | 0.037 | 0.027 | 0.015 | 0.017 |
| Left Parietal Operculum Cortex | 0.812 | 0.750 | 0.029 | 0.028 | 0.019 | 0.018 | 0.024 | 0.024 | 0.014 | 0.018 |
| Right Parietal Operculum Cortex | 1.072 | 0.988 | 0.030 | 0.028 | 0.021 | 0.019 | 0.031 | 0.027 | 0.015 | 0.018 |
| Left Planum Polare | 0.923 | 0.866 | 0.031 | 0.026 | 0.022 | 0.019 | 0.037 | 0.028 | 0.016 | 0.017 |
| Right Planum Polare | 1.339 | 1.024 | 0.029 | 0.024 | 0.021 | 0.018 | 0.036 | 0.026 | 0.016 | 0.017 |
| Left Heschl's Gyrus (includes H1 and H2) | 0.971 | 0.901 | 0.028 | 0.027 | 0.021 | 0.018 | 0.025 | 0.022 | 0.017 | 0.022 |
| Right Heschl's Gyrus (includes H1 and H2) | 0.917 | 0.841 | 0.024 | 0.025 | 0.017 | 0.016 | 0.026 | 0.024 | 0.014 | 0.021 |
| Left Planum Temporale | 0.994 | 0.912 | 0.024 | 0.022 | 0.017 | 0.016 | 0.032 | 0.026 | 0.014 | 0.020 |
| Right Planum Temporale | 1.429 | 1.176 | 0.026 | 0.027 | 0.017 | 0.016 | 0.023 | 0.022 | 0.014 | 0.020 |
| Left Supracalcarine Cortex | 1.340 | 1.098 | 0.027 | 0.026 | 0.019 | 0.018 | 0.030 | 0.027 | 0.013 | 0.018 |
| Right Supracalcarine Cortex | 1.135 | 0.953 | 0.024 | 0.023 | 0.018 | 0.016 | 0.030 | 0.026 | 0.013 | 0.019 |
| Left Occipital Pole | 1.180 | 0.954 | 0.030 | 0.023 | 0.021 | 0.018 | 0.042 | 0.026 | 0.014 | 0.016 |
| Right Occipital Pole | 1.284 | 1.025 | 0.028 | 0.022 | 0.019 | 0.017 | 0.036 | 0.023 | 0.014 | 0.015 |

**Supplementary Table 2:** Table showing the F-values for main effect of age group for BOLD signal variability ($SD_{BOLD}$). Statistical significance was determined using nonparametric ANCOVAs corrected for multiple comparisons by false discovery rates (FDR; Benjamini and Hochberg, 1995). The nonparametric ANCOVAs with $SD_{BOLD}$ as dependent variable demonstrated that there was a significant main effect of age group in 72 ROIs, while no significant effect of sex was observed.

| **ROI** | **F-value: age** |
|---|---|
| Left.Angular.Gyrus | 20.9868 |
| Left.Central.Opercular.Cortex | 15.7773 |
| Left.Cingulate.Gyrus..anterior.division | 37.6699 |
| Left.Cingulate.Gyrus..posterior.division | 24.6328 |
| Left.Cuneal.Cortex | 20.4775 |
| Left.Frontal.Medial.Cortex | 16.5504 |
| Left.Frontal.Orbital.Cortex | 59.1622 |
| Left.Frontal.Pole | 27.4942 |
| Left.Heschls.Gyrus..includes.H1.and.H2. | 15.867 |
| Left.Inferior.Frontal.Gyrus..pars.opercularis | 42.0627 |
| Left.Inferior.Frontal.Gyrus..pars.triangularis | 17.0069 |
| Left.Inferior.Temporal.Gyrus..posterior.division | 28.2274 |
| Left.Insular.Cortex | 38.1285 |
| Left.Intracalcarine.Cortex | 34.2193 |
| Left.Juxtapositional.Lobule.Cortex..formerly.Supplementary.Motor.Cortex. | 40.0826 |
| Left.Middle.Frontal.Gyrus | 35.6619 |
| Left.Middle.Temporal.Gyrus..temporooccipital.part | 24.0403 |
| Left.Occipital.Fusiform.Gyrus | 35.3288 |
| Left.Occipital.Pole | 43.1077 |
| Left.Paracingulate.Gyrus | 54.3683 |
| Left.Parahippocampal.Gyrus..anterior.division | 60.2925 |
| Left.Parahippocampal.Gyrus..posterior.division | 38.4558 |

| | |
|---|---|
| Left.Parietal.Operculum.Cortex | 25.0314 |
| Left.Postcentral.Gyrus | 29.5352 |
| Left.Precuneous.Cortex | 28.1132 |
| Left.Subcallosal.Cortex | 35.3522 |
| Left.Superior.Frontal.Gyrus | 16.076 |
| Left.Superior.Temporal.Gyrus..posterior.division | 19.4259 |
| Left.Supracalcarine.Cortex | 25.8525 |
| Left.Supramarginal.Gyrus..anterior.division | 15.3458 |
| Left.Temporal.Fusiform.Cortex..anterior.division | 17.3434 |
| Left.Temporal.Fusiform.Cortex..posterior.division | 37.6047 |
| Left.Temporal.Occipital.Fusiform.Cortex | 41.0751 |
| Right.Angular.Gyrus | 22.823 |
| Right.Cingulate.Gyrus..anterior.division | 35.6207 |
| Right.Cingulate.Gyrus..posterior.division | 28.4509 |
| Right.Cuneal.Cortex | 25.6256 |
| Right.Frontal.Medial.Cortex | 44.9473 |
| Right.Frontal.Orbital.Cortex | 61.1423 |
| Right.Frontal.Pole | 36.7283 |
| Right.Inferior.Frontal.Gyrus..pars.opercularis | 14.9685 |
| Right.Inferior.Frontal.Gyrus..pars.triangularis | 15.799 |
| Right.Inferior.Temporal.Gyrus..anterior.division | 21.551 |
| Right.Inferior.Temporal.Gyrus..posterior.division | 15.0549 |
| Right.Insular.Cortex | 18.3847 |
| Right.Intracalcarine.Cortex | 13.3254 |
| Right.Juxtapositional.Lobule.Cortex..formerly.Supplementary.Motor.Cortex. | 36.4605 |
| Right.Lateral.Occipital.Cortex..inferior.division | 33.6556 |
| Right.Lateral.Occipital.Cortex..superior.division | 21.5155 |
| Right.Lingual.Gyrus | 25.0696 |

| | |
|---|---|
| Right.Middle.Temporal.Gyrus..anterior.division | 42.8831 |
| Right.Occipital.Fusiform.Gyrus | 31.6213 |
| Right.Occipital.Pole | 31.9927 |
| Right.Paracingulate.Gyrus | 34.8449 |
| Right.Parahippocampal.Gyrus..anterior.division | 37.5431 |
| Right.Parietal.Operculum.Cortex | 18.2991 |
| Right.Planum.Polare | 57.0714 |
| Right.Planum.Temporale | 25.7173 |
| Right.Postcentral.Gyrus | 20.1925 |
| Right.Precentral.Gyrus | 18.1681 |
| Right.Precuneous.Cortex | 24.8254 |
| Right.Subcallosal.Cortex | 41.1512 |
| Right.Superior.Frontal.Gyrus | 25.8665 |
| Right.Superior.Parietal.Lobule | 23.6064 |
| Right.Superior.Temporal.Gyrus..anterior.division | 17.9951 |
| Right.Superior.Temporal.Gyrus..posterior.division | 18.4805 |
| Right.Supracalcarine.Cortex | 28.7788 |
| Right.Supramarginal.Gyrus..anterior.division | 18.6616 |
| Right.Supramarginal.Gyrus..posterior.division | 23.6768 |
| Right.Temporal.Fusiform.Cortex..anterior.division | 23.371 |
| Right.Temporal.Fusiform.Cortex..posterior.division | 24.4233 |
| Right.Temporal.Occipital.Fusiform.Cortex | 31.6812 |

**Supplementary Table 3:** Table showing the F-values for the main effect of age group for EEG signal variability (1-3 Hz, $SD_{DELTA}$). Statistical significance was determined using nonparametric ANCOVAs corrected for multiple comparisons by false discovery rates (FDR; Benjamini and Hochberg, 1995). The nonparametric ANCOVAs with $SD_{DELTA}$ as dependent variable demonstrated that there was a significant main effect of age group in 14 ROIs, and sex in in 20 ROIs.

| ROI | F-value: age | ROI | F-value: sex |
|---|---|---|---|
| Left.Cingulate.Gyrus..posterior.division | 17.041 | Left.Cuneal.Cortex | 15.463 |
| Left.Cuneal.Cortex | 20.8528 | Left.Intracalcarine.Cortex | 20.018 |
| Left.Intracalcarine.Cortex | 12.579 | Left.Lingual.Gyrus | 13.828 |
| Left.Juxtapositional.Lobule.Cortex..formerly.Supplementary.Motor.Cortex. | 13.1327 | Left.Parietal.Operculum.Cortex | 13.243 |
| Left.Lateral.Occipital.Cortex..superior.division | 15.2747 | Left.Supracalcarine.Cortex | 21.285 |
| Left.Precuneous.Cortex | 19.2268 | Right.Cuneal.Cortex | 17.36 |
| Left.Supracalcarine.Cortex | 16.815 | Right.Heschls.Gyrus..includes.H1.and.H2. | 15.627 |
| Right.Cingulate.Gyrus..posterior.division | 17.85 | Right.Inferior.Temporal.Gyrus..posterior.division | 16.731 |
| Right.Cuneal.Cortex | 20.1038 | Right.Inferior.Temporal.Gyrus..temporooccipital.part | 26.594 |
| Right.Intracalcarine.Cortex | 15.1102 | Right.Intracalcarine.Cortex | 26.625 |
| Right.Juxtapositional.Lobule.Cortex..formerly.Supplementary.Motor.Cortex. | 13.7918 | Right.Lateral.Occipital.Cortex..inferior.division | 24.394 |
| Right.Lateral.Occipital.Cortex..superior.division | 19.2019 | Right.Lingual.Gyrus | 19.964 |
| Right.Precuneous.Cortex | 20.9403 | Right.Middle.Temporal.Gyrus..temporooccipital.part | 24.628 |
| Right.Supracalcarine.Cortex | 17.4001 | Right.Occipital.Fusiform.Gyrus | 19.042 |
| | | Right.Occipital.Pole | 14.198 |
| | | Right.Parahippocampal.Gyrus..posterior.division | 15.023 |
| | | Right.Parietal.Operculum.Cortex | 15.099 |
| | | Right.Planum.Temporale | 16.428 |
| | | Right.Supracalcarine.Cortex | 24.14 |
| | | Right.Temporal.Fusiform.Cortex..posterior.division | 16.515 |
| | | Right.Temporal.Occipital.Fusiform.Cortex | 23.004 |

**Supplementary Table 4:** Table showing the F-values for the main effect of age group for EEG signal variability (4-7 Hz, $SD_{THETA}$). Statistical significance was determined using nonparametric ANCOVAs corrected for multiple comparisons by false discovery rates (FDR; Benjamini and Hochberg, 1995). The nonparametric ANCOVAs with $SD_{THETA}$ as dependent variable demonstrated that there was a significant main effect of age group in 16 ROIs, and sex in in 74 ROIs.

| ROI | F-value: age | ROI | F-value: sex |
|---|---|---|---|
| Left.Cingulate.Gyrus..anterior.division | 33.5753 | Left.Angular.Gyrus | 18.207 |
| Left.Cingulate.Gyrus..posterior.division | 19.2188 | Left.Central.Opercular.Cortex | 13.609 |
| Left.Juxtapositional.Lobule.Cortex..formerly.Supplementary.Motor.Cortex. | 39.3985 | Left.Cingulate.Gyrus..anterior.division | 13.504 |
| Left.Middle.Frontal.Gyrus | 24.3858 | Left.Cingulate.Gyrus..posterior.division | 17.748 |
| Left.Paracingulate.Gyrus | 22.3193 | Left.Cuneal.Cortex | 22.505 |
| Left.Precentral.Gyrus | 17.0336 | Left.Frontal.Orbital.Cortex | 13.146 |
| Left.Precuneous.Cortex | 13.1632 | Left.Frontal.Pole | 16.278 |
| Left.Superior.Frontal.Gyrus | 33.9343 | Left.Inferior.Frontal.Gyrus..pars.opercularis | 12.817 |
| Right.Cingulate.Gyrus..anterior.division | 32.1706 | Left.Inferior.Temporal.Gyrus..posterior.division | 13.732 |
| Right.Cingulate.Gyrus..posterior.division | 18.9454 | Left.Inferior.Temporal.Gyrus..temporooccipital.part | 16.711 |
| Right.Juxtapositional.Lobule.Cortex..formerly.Supplementary.Motor.Cortex. | 40.3098 | Left.Intracalcarine.Cortex | 27.158 |
| Right.Middle.Frontal.Gyrus | 23.3526 | Left.Juxtapositional.Lobule.Cortex..formerly.Supplementary.Motor.Cortex. | 13.129 |
| Right.Paracingulate.Gyrus | 19.4883 | Left.Lateral.Occipital.Cortex..inferior.division | 16.541 |
| Right.Precentral.Gyrus | 17.9796 | Left.Lateral.Occipital.Cortex..superior.division | 13.959 |
| Right.Superior.Frontal.Gyrus | 33.4244 | Left.Lingual.Gyrus | 22.478 |
| Right.Superior.Parietal.Lobule | 13.5708 | Left.Middle.Temporal.Gyrus..temporooccipital.part | 16.103 |
| | | Left.Occipital.Fusiform.Gyrus | 18.143 |
| | | Left.Occipital.Pole | 18.184 |
| | | Left.Parahippocampal.Gyrus..posterior.division | 14.667 |

| | |
|---|---|
| Left.Parietal.Operculum.Cortex | 21.209 |
| Left.Planum.Temporale | 18.513 |
| Left.Postcentral.Gyrus | 16.801 |
| Left.Precentral.Gyrus | 16.951 |
| Left.Precuneous.Cortex | 17.475 |
| Left.Subcallosal.Cortex | 13.415 |
| Left.Supracalcarine.Cortex | 27.108 |
| Left.Supramarginal.Gyrus..anterior.division | 22.636 |
| Left.Supramarginal.Gyrus..posterior.division | 18.42 |
| Left.Temporal.Fusiform.Cortex..posterior.division | 12.846 |
| Left.Temporal.Occipital.Fusiform.Cortex | 18.695 |
| Right.Angular.Gyrus | 17.082 |
| Right.Central.Opercular.Cortex | 23.027 |
| Right.Cingulate.Gyrus..anterior.division | 13.094 |
| Right.Cingulate.Gyrus..posterior.division | 17.068 |
| Right.Cuneal.Cortex | 22.446 |
| Right.Frontal.Operculum.Cortex | 25.517 |
| Right.Frontal.Orbital.Cortex | 17.066 |
| Right.Frontal.Pole | 12.686 |
| Right.Heschls.Gyrus..includes.H1.and.H2. | 16.19 |
| Right.Inferior.Frontal.Gyrus..pars.opercularis | 25.941 |
| Right.Inferior.Frontal.Gyrus..pars.triangularis | 24.437 |
| Right.Inferior.Temporal.Gyrus..anterior.division | 19.63 |
| Right.Inferior.Temporal.Gyrus..posterior.division | 22.865 |
| Right.Inferior.Temporal.Gyrus..temporooccipital.part | 28.425 |
| Right.Insular.Cortex | 22.486 |
| Right.Intracalcarine.Cortex | 29.957 |
| Right.Juxtapositional.Lobule.Cortex..formerly.Supplementary.Motor | 13.499 |

| | |
|---|---|
| Right.Lateral.Occipital.Cortex..inferior.division | 30.063 |
| Right.Lateral.Occipital.Cortex..superior.division | 13.899 |
| Right.Lingual.Gyrus | 25.891 |
| Right.Middle.Frontal.Gyrus | 18.676 |
| Right.Middle.Temporal.Gyrus..anterior.division | 13.832 |
| Right.Middle.Temporal.Gyrus..posterior.division | 21.779 |
| Right.Middle.Temporal.Gyrus..temporooccipital.part | 28.874 |
| Right.Occipital.Fusiform.Gyrus | 27.265 |
| Right.Occipital.Pole | 28.335 |
| Right.Parahippocampal.Gyrus..anterior.division | 15.973 |
| Right.Parahippocampal.Gyrus..posterior.division | 17.709 |
| Right.Parietal.Operculum.Cortex | 16.92 |
| Right.Planum.Polare | 17.124 |
| Right.Planum.Temporale | 17.351 |
| Right.Postcentral.Gyrus | 18.97 |
| Right.Precentral.Gyrus | 21.799 |
| Right.Precuneous.Cortex | 16.222 |
| Right.Subcallosal.Cortex | 14.188 |
| Right.Superior.Temporal.Gyrus..anterior.division | 13.072 |
| Right.Superior.Temporal.Gyrus..posterior.division | 19.288 |
| Right.Supracalcarine.Cortex | 27.779 |
| Right.Supramarginal.Gyrus..anterior.division | 19.774 |
| Right.Supramarginal.Gyrus..posterior.division | 15.411 |
| Right.Temporal.Fusiform.Cortex..anterior.division | 18.158 |
| Right.Temporal.Fusiform.Cortex..posterior.division | 21.228 |
| Right.Temporal.Occipital.Fusiform.Cortex | 26.317 |
| Right.Temporal.Pole | 16.482 |

**Supplementary Table 5:** Table showing the F-values for the main effect of age group for EEG signal variability (8-12 Hz, $SD_{ALPHA}$). Statistical significance was determined using nonparametric ANCOVAs corrected for multiple comparisons by false discovery rates (FDR; Benjamini and Hochberg, 1995). The nonparametric ANCOVAs with $SD_{ALPHA}$ as dependent variable demonstrated that there was a significant main effect of age group in 20 ROIs, and sex in in 4 ROIs.

| **ROI** | **F-value: age** | **ROI** | **F-value: sex** |
|---|---|---|---|
| Left.Cingulate.Gyrus..anterior.division | 14.2391 | Left.Frontal.Pole | 16.512 |
| Left.Cingulate.Gyrus..posterior.division | 18.9122 | Right.Frontal.Orbital.Cortex | 13.217 |
| Left.Cuneal.Cortex | 19.5004 | Right.Inferior.Frontal.Gyrus..pars.triangularis | 14.278 |
| Left.Intracalcarine.Cortex | 13.9943 | Right.Temporal.Pole | 12.887 |
| Left.Juxtapositional.Lobule.Cortex..formerly.Supplementary.Motor.Cortex. | 18.2498 | | |
| Left.Lateral.Occipital.Cortex..superior.division | 14.8073 | | |
| Left.Occipital.Pole | 13.6297 | | |
| Left.Precuneous.Cortex | 18.1658 | | |
| Left.Supracalcarine.Cortex | 18.7095 | | |
| Right.Cingulate.Gyrus..anterior.division | 14.9683 | | |
| Right.Cingulate.Gyrus..posterior.division | 19.3745 | | |
| Right.Cuneal.Cortex | 20.1265 | | |
| Right.Intracalcarine.Cortex | 17.8042 | | |
| Right.Juxtapositional.Lobule.Cortex..formerly.Supplementary.Motor.Cortex. | 18.9006 | | |
| Right.Lateral.Occipital.Cortex..superior.division | 18.2382 | | |
| Right.Occipital.Pole | 17.1219 | | |
| Right.Precuneous.Cortex | 19.3677 | | |
| Right.Superior.Frontal.Gyrus | 14.4975 | | |
| Right.Superior.Parietal.Lobule | 13.8287 | | |
| Right.Supracalcarine.Cortex | 19.6116 | | |

**Supplementary Table 6:** Table showing the F-values for the main effect of age group for EEG signal variability (15-25 Hz, $SD_{BETA}$). Statistical significance was determined using nonparametric ANCOVAs corrected for multiple comparisons by false discovery rates (FDR; Benjamini and Hochberg, 1995). The nonparametric ANCOVA analyses with $SD_{BETA}$ as dependent variable demonstrated that there was a significant main effect of age group in 19 ROIs, and sex in 69 ROIs.

| ROI | F-value: age | ROI | F-value: sex |
|---|---|---|---|
| Left.Central.Opercular.Cortex | 16.5945 | Left.Angular.Gyrus | 23.069 |
| Left.Heschls.Gyrus..includes.H1.and.H2. | 20.3049 | Left.Cingulate.Gyrus..anterior.division | 12.198 |
| Left.Inferior.Temporal.Gyrus..posterior.division | 15.6087 | Left.Cingulate.Gyrus..posterior.division | 23.953 |
| Left.Insular.Cortex | 12.903 | Left.Cuneal.Cortex | 29.101 |
| Left.Middle.Temporal.Gyrus..posterior.division | 13.2449 | Left.Frontal.Medial.Cortex | 20.926 |
| Left.Parietal.Operculum.Cortex | 21.6143 | Left.Frontal.Operculum.Cortex | 12.549 |
| Left.Planum.Temporale | 21.1029 | Left.Frontal.Orbital.Cortex | 22.762 |
| Left.Superior.Temporal.Gyrus..posterior.division | 14.4852 | Left.Frontal.Pole | 19.186 |
| Left.Supramarginal.Gyrus..anterior.division | 19.6781 | Left.Heschls.Gyrus..includes.H1.and.H2. | 18.878 |
| Left.Supramarginal.Gyrus..posterior.division | 15.7818 | Left.Inferior.Temporal.Gyrus..posterior.division | 14.641 |
| Right.Central.Opercular.Cortex | 17.2968 | Left.Inferior.Temporal.Gyrus..temporooccipital.part | 24.317 |
| Right.Heschls.Gyrus..includes.H1.and.H2. | 13.1379 | Left.Insular.Cortex | 15.927 |
| Right.Middle.Temporal.Gyrus..anterior.division | 15.6176 | Left.Intracalcarine.Cortex | 35.249 |
| Right.Middle.Temporal.Gyrus..posterior.division | 15.253 | Left.Lateral.Occipital.Cortex..inferior.division | 22.746 |
| Right.Parietal.Operculum.Cortex | 12.510 | Left.Lateral.Occipital.Cortex..superior.division | 19.834 |
| Right.Planum.Polare | 14.949 | Left.Lingual.Gyrus | 32.404 |
| Right.Planum.Temporale | 12.617 | Left.Middle.Temporal.Gyrus..temporooccipital.part | 17.782 |
| Right.Superior.Temporal.Gyrus..anterior.division | 19.6635 | Left.Occipital.Fusiform.Gyrus | 24.141 |
| Right.Superior.Temporal.Gyrus..posterior.division | 18.8534 | Left.Occipital.Pole | 23.17 |
| | | Paracingulate.Gyrus | 14.818 |
| | | Left.Parahippocampal.Gyrus..anterior.division | 23.933 |

| | |
|---|---|
| Left.Parahippocampal.Gyrus..posterior.division | 30.177 |
| Left.Parietal.Operculum.Cortex | 19.769 |
| Left.Planum.Temporale | 20.791 |
| Left.Precuneous.Cortex | 25.556 |
| Left.Subcallosal.Cortex | 22.89 |
| Left.Superior.Parietal.Lobule | 13.534 |
| Left.Supracalcarine.Cortex | 34.165 |
| Left.Supramarginal.Gyrus..posterior.division | 19.854 |
| Left.Temporal.Fusiform.Cortex..anterior.division | 17.146 |
| Left.Temporal.Fusiform.Cortex..posterior.division | 25.091 |
| Left.Temporal.Occipital.Fusiform.Cortex | 30.718 |
| Left.Temporal.Pole | 15.41 |
| Right.Angular.Gyrus | 26.346 |
| Right.Central.Opercular.Cortex | 13.78 |
| Right.Cingulate.Gyrus..posterior.division | 23.764 |
| Right.Cuneal.Cortex | 26.278 |
| Right.Frontal.Medial.Cortex | 16.911 |
| Right.Frontal.Orbital.Cortex | 13.877 |
| Right.Heschls.Gyrus..includes.H1.and.H2. | 16.353 |
| Right.Inferior.Temporal.Gyrus..anterior.division | 15.577 |
| Right.Inferior.Temporal.Gyrus..posterior.division | 18.262 |
| Right.Inferior.Temporal.Gyrus..temporooccipital.part | 29.167 |
| Right.Insular.Cortex | 16.383 |
| Right.Intracalcarine.Cortex | 30.856 |
| Right.Lateral.Occipital.Cortex..inferior.division | 35.716 |
| Right.Lateral.Occipital.Cortex..superior.division | 19.643 |
| Right.Lingual.Gyrus | 27.537 |
| Right.Middle.Temporal.Gyrus..posterior.division | 16.15 |

| | |
|---|---|
| Right.Middle.Temporal.Gyrus..temporooccipital.part | 32.438 |
| Right.Occipital.Fusiform.Gyrus | 24.567 |
| Right.Occipital.Pole | 30.231 |
| Right.Parahippocampal.Gyrus..anterior.division | 20.351 |
| Right.Parahippocampal.Gyrus..posterior.division | 25.145 |
| Right.Parietal.Operculum.Cortex | 21.039 |
| Right.Planum.Polare | 14.552 |
| Right.Planum.Temporale | 21.152 |
| Right.Postcentral.Gyrus | 13.107 |
| Right.Precuneous.Cortex | 22.867 |
| Right.Subcallosal.Cortex | 18.68 |
| Right.Superior.Parietal.Lobule | 13.68 |
| Right.Superior.Temporal.Gyrus..posterior.division | 14.929 |
| Right.Supracalcarine.Cortex | 30.665 |
| Right.Supramarginal.Gyrus..anterior.division | 15.081 |
| Right.Supramarginal.Gyrus..posterior.division | 22.485 |
| Right.Temporal.Fusiform.Cortex..anterior.division | 17.436 |
| Right.Temporal.Fusiform.Cortex..posterior.division | 21.616 |
| Right.Temporal.Occipital.Fusiform.Cortex | 28.231 |
| Right.Temporal.Pole | 14.486 |

**Supplementary Table 7.** Spearman correlation of $SD_{BOLD}$ with $SD_{EEG}$ for each frequency band in young subjects (N=135). None of the pairwise correlations between $SD_{BOLD}$ and $SD_{EEG}$ were statistically significant.

| ROI | rho $SD_{DELTA}$ | rho $SD_{THETA}$ | rho $SD_{ALPHA}$ | rho $SD_{BETA}$ |
|---|---|---|---|---|
| Left Angular Gyrus | 0.005 | 0.015 | 0.014 | 0.071 |
| Left Central Opercular Cortex | 0.037 | 0.001 | 0.076 | 0.034 |
| Left Cingulate Gyrus, anterior division | -0.090 | -0.077 | 0.091 | -0.028 |
| Left Cingulate Gyrus, posterior division | -0.096 | -0.048 | -0.018 | 0.042 |
| Left Cuneal Cortex | -0.166 | -0.153 | -0.040 | -0.055 |
| Left Frontal Medial Cortex | -0.009 | -0.083 | -0.105 | -0.100 |
| Left Frontal Operculum Cortex | -0.067 | -0.128 | 0.074 | -0.094 |
| Left Frontal Orbital Cortex | 0.035 | -0.010 | 0.137 | 0.107 |
| Left Frontal Pole | 0.110 | -0.018 | -0.029 | -0.052 |
| Left Heschl's Gyrus (includes H1 and H2) | -0.019 | 0.029 | 0.112 | -0.096 |
| Left Inferior Frontal Gyrus, pars opercularis | 0.040 | -0.082 | 0.063 | -0.091 |
| Left Inferior Frontal Gyrus, pars triangularis | 0.031 | -0.114 | 0.064 | -0.041 |
| Left Inferior Temporal Gyrus, anterior division | 0.035 | 0.023 | 0.099 | 0.012 |
| Left Inferior Temporal Gyrus, posterior division | 0.047 | 0.078 | 0.116 | 0.034 |
| Left Inferior Temporal Gyrus, temporooccipital part | 0.030 | -0.002 | 0.025 | -0.089 |
| Left Insular Cortex | 0.023 | -0.134 | -0.017 | -0.077 |
| Left Intracalcarine Cortex | 0.018 | 0.028 | 0.072 | 0.032 |
| Left Juxtapositional Lobule Cortex (formerly Supplementary Motor Cortex) | -0.050 | -0.099 | 0.077 | 0.022 |
| Left Lateral Occipital Cortex, inferior division | -0.048 | -0.030 | -0.078 | -0.077 |
| Left Lateral Occipital Cortex, superior division | -0.062 | -0.081 | 0.018 | -0.088 |
| Left Lingual Gyrus | 0.131 | 0.060 | -0.022 | -0.035 |
| Left Middle Frontal Gyrus | 0.041 | -0.066 | 0.131 | 0.015 |
| Left Middle Temporal Gyrus, anterior division | 0.004 | -0.063 | -0.014 | -0.173 |
| Left Middle Temporal Gyrus, posterior division | 0.115 | 0.005 | -0.041 | -0.029 |
| Left Middle Temporal Gyrus, temporooccipital part | 0.072 | 0.037 | 0.071 | -0.092 |
| Left Occipital Fusiform Gyrus | 0.146 | 0.149 | 0.115 | 0.026 |
| Left Occipital Pole | -0.017 | 0.052 | 0.036 | 0.035 |
| Left Paracingulate Gyrus | -0.044 | -0.087 | -0.012 | -0.036 |
| Left Parahippocampal Gyrus, anterior division | 0.024 | 0.021 | 0.121 | 0.027 |
| Left Parahippocampal Gyrus, posterior division | -0.086 | 0.030 | 0.117 | 0.162 |
| Left Parietal Operculum Cortex | -0.130 | -0.064 | 0.039 | -0.110 |
| Left Planum Polare | 0.030 | 0.018 | 0.073 | -0.004 |
| Left Planum Temporale | -0.066 | 0.009 | 0.120 | -0.071 |
| Left Postcentral Gyrus | -0.019 | -0.032 | 0.139 | -0.060 |
| Left Precentral Gyrus | -0.015 | -0.074 | 0.091 | -0.056 |
| Left Precuneous Cortex | -0.029 | -0.070 | 0.107 | -0.044 |
| Left Subcallosal Cortex | -0.038 | -0.087 | 0.034 | -0.074 |
| Left Superior Frontal Gyrus | -0.108 | -0.139 | 0.027 | -0.038 |

| | | | | |
|---|---|---|---|---|
| Left Superior Parietal Lobule | 0.087 | 0.041 | 0.135 | -0.084 |
| Left Superior Temporal Gyrus, anterior division | -0.010 | -0.074 | 0.033 | 0.064 |
| Left Superior Temporal Gyrus, posterior division | -0.059 | -0.045 | -0.047 | -0.087 |
| Left Supracalcarine Cortex | -0.076 | -0.071 | 0.036 | 0.016 |
| Left Supramarginal Gyrus, anterior division | 0.026 | -0.060 | 0.001 | -0.057 |
| Left Supramarginal Gyrus, posterior division | -0.005 | 0.066 | 0.106 | 0.043 |
| Left Temporal Fusiform Cortex, anterior division | 0.188 | 0.120 | 0.051 | 0.031 |
| Left Temporal Fusiform Cortex, posterior division | 0.056 | 0.052 | 0.144 | 0.075 |
| Left Temporal Occipital Fusiform Cortex | 0.096 | 0.094 | 0.128 | 0.092 |
| Left Temporal Pole | 0.224 | 0.105 | 0.152 | -0.010 |
| Right Angular Gyrus | -0.010 | 0.025 | 0.010 | 0.046 |
| Right Central Opercular Cortex | 0.084 | -0.015 | 0.068 | -0.030 |
| Right Cingulate Gyrus, anterior division | -0.079 | -0.085 | 0.062 | -0.046 |
| Right Cingulate Gyrus, posterior division | -0.090 | -0.048 | -0.004 | 0.021 |
| Right Cuneal Cortex | -0.118 | -0.096 | 0.017 | -0.074 |
| Right Frontal Medial Cortex | 0.042 | -0.043 | 0.063 | -0.050 |
| Right Frontal Operculum Cortex | 0.004 | -0.017 | 0.089 | -0.075 |
| Right Frontal Orbital Cortex | 0.085 | -0.024 | 0.100 | -0.032 |
| Right Frontal Pole | 0.149 | -0.022 | 0.014 | -0.061 |
| Right Heschl's Gyrus (includes H1 and H2) | -0.085 | -0.045 | 0.065 | -0.071 |
| Right Inferior Frontal Gyrus, pars opercularis | -0.020 | -0.029 | 0.127 | -0.138 |
| Right Inferior Frontal Gyrus, pars triangularis | -0.047 | -0.092 | 0.061 | -0.170 |
| Right Inferior Temporal Gyrus, anterior division | 0.013 | -0.018 | 0.058 | 0.028 |
| Right Inferior Temporal Gyrus, posterior division | 0.179 | 0.070 | 0.128 | -0.048 |
| Right Inferior Temporal Gyrus, temporooccipital part | 0.106 | 0.068 | 0.177 | 0.093 |
| Right Insular Cortex | -0.034 | -0.087 | 0.043 | -0.054 |
| Right Intracalcarine Cortex | -0.063 | -0.121 | 0.008 | -0.065 |
| Right Juxtapositional Lobule Cortex (formerly Supplementary Motor Cortex) | -0.040 | -0.142 | 0.015 | -0.049 |
| Right Lateral Occipital Cortex, inferior division | -0.025 | 0.005 | 0.081 | 0.026 |
| Right Lateral Occipital Cortex, superior division | -0.008 | -0.037 | 0.044 | -0.113 |
| Right Lingual Gyrus | 0.040 | 0.033 | 0.058 | 0.034 |
| Right Middle Frontal Gyrus | -0.167 | -0.083 | 0.063 | -0.143 |
| Right Middle Temporal Gyrus, anterior division | 0.135 | 0.049 | 0.054 | -0.113 |
| Right Middle Temporal Gyrus, posterior division | 0.002 | 0.080 | 0.191 | 0.060 |
| Right Middle Temporal Gyrus, temporooccipital part | 0.026 | 0.011 | 0.094 | 0.049 |
| Right Occipital Fusiform Gyrus | 0.095 | 0.039 | 0.022 | 0.007 |
| Right Occipital Pole | 0.019 | -0.050 | 0.025 | -0.036 |
| Right Paracingulate Gyrus | -0.063 | -0.041 | 0.034 | -0.012 |
| Right Parahippocampal Gyrus, anterior division | -0.027 | -0.005 | 0.071 | 0.102 |
| Right Parahippocampal Gyrus, posterior division | 0.020 | -0.037 | 0.037 | 0.024 |
| Right Parietal Operculum Cortex | -0.066 | -0.039 | 0.124 | -0.075 |

| | | | | |
|---|---|---|---|---|
| Right Planum Polare | 0.186 | 0.073 | 0.105 | -0.021 |
| Right Planum Temporale | -0.028 | -0.043 | 0.074 | 0.070 |
| Right Postcentral Gyrus | 0.070 | 0.026 | 0.133 | -0.056 |
| Right Precentral Gyrus | -0.031 | -0.047 | 0.134 | -0.040 |
| Right Precuneous Cortex | -0.099 | -0.145 | 0.053 | -0.200 |
| Right Subcallosal Cortex | -0.084 | -0.086 | 0.098 | 0.054 |
| Right Superior Frontal Gyrus | -0.100 | -0.158 | -0.026 | -0.116 |
| Right Superior Parietal Lobule | -0.035 | -0.067 | 0.080 | -0.103 |
| Right Superior Temporal Gyrus, anterior division | 0.144 | 0.056 | 0.050 | 0.021 |
| Right Superior Temporal Gyrus, posterior division | 0.067 | 0.068 | 0.064 | 0.004 |
| Right Supracalcarine Cortex | -0.004 | 0.049 | 0.089 | 0.065 |
| Right Supramarginal Gyrus, anterior division | 0.134 | 0.092 | 0.116 | 0.046 |
| Right Supramarginal Gyrus, posterior division | 0.058 | 0.055 | 0.096 | 0.052 |
| Right Temporal Fusiform Cortex, anterior division | 0.038 | -0.019 | 0.013 | 0.142 |
| Right Temporal Fusiform Cortex, posterior division | 0.012 | 0.011 | 0.075 | 0.103 |
| Right Temporal Occipital Fusiform Cortex | -0.013 | 0.018 | 0.115 | 0.098 |
| Right Temporal Pole | 0.135 | 0.006 | 0.044 | -0.129 |

*$p_{FDR} < 0.05$; **$p_{FDR} < 0.01$; ***$p_{FDR} < 0.001$, 2-tailed*

**Supplementary Table 8.** Spearman correlation of $SD_{BOLD}$ with $SD_{EEG}$ for each frequency band in old subjects (N=54). None of the pairwise correlations between $SD_{BOLD}$ and $SD_{EEG}$ were statistically significant.

| ROI | rho $SD_{DELTA}$ | rho $SD_{THETA}$ | rho $SD_{ALPHA}$ | rho $SD_{BETA}$ |
|---|---|---|---|---|
| Left Angular Gyrus | -0.118 | -0.100 | -0.001 | -0.192 |
| Left Central Opercular Cortex | 0.129 | 0.132 | -0.006 | 0.204 |
| Left Cingulate Gyrus, anterior division | -0.043 | 0.175 | -0.034 | 0.010 |
| Left Cingulate Gyrus, posterior division | 0.014 | -0.077 | 0.012 | -0.301 |
| Left Cuneal Cortex | -0.134 | -0.157 | -0.096 | -0.387 |
| Left Frontal Medial Cortex | -0.165 | -0.129 | -0.265 | 0.061 |
| Left Frontal Operculum Cortex | 0.173 | 0.168 | 0.063 | 0.029 |
| Left Frontal Orbital Cortex | 0.085 | 0.192 | 0.020 | 0.021 |
| Left Frontal Pole | 0.038 | -0.009 | -0.044 | -0.080 |
| Left Heschl's Gyrus (includes H1 and H2) | -0.066 | -0.152 | -0.166 | 0.027 |
| Left Inferior Frontal Gyrus, pars opercularis | 0.039 | 0.041 | -0.039 | 0.177 |
| Left Inferior Frontal Gyrus, pars triangularis | 0.107 | 0.113 | 0.033 | 0.086 |
| Left Inferior Temporal Gyrus, anterior division | 0.111 | 0.153 | 0.088 | 0.278 |
| Left Inferior Temporal Gyrus, posterior division | 0.072 | 0.040 | -0.035 | 0.040 |
| Left Inferior Temporal Gyrus, temporooccipital part | 0.016 | 0.074 | 0.034 | -0.066 |
| Left Insular Cortex | 0.225 | 0.042 | -0.079 | 0.026 |
| Left Intracalcarine Cortex | 0.102 | 0.130 | 0.254 | 0.172 |
| Left Juxtapositional Lobule Cortex (formerly Supplementary Motor Cortex) | -0.054 | -0.175 | -0.035 | -0.059 |
| Left Lateral Occipital Cortex, inferior division | 0.036 | -0.027 | -0.057 | -0.184 |
| Left Lateral Occipital Cortex, superior division | 0.130 | 0.033 | 0.013 | -0.083 |
| Left Lingual Gyrus | -0.181 | -0.101 | -0.044 | -0.130 |
| Left Middle Frontal Gyrus | 0.159 | 0.161 | -0.009 | -0.008 |
| Left Middle Temporal Gyrus, anterior division | -0.004 | 0.000 | -0.012 | 0.349 |
| Left Middle Temporal Gyrus, posterior division | -0.007 | -0.073 | -0.197 | 0.077 |
| Left Middle Temporal Gyrus, temporooccipital part | 0.042 | 0.203 | 0.128 | 0.015 |
| Left Occipital Fusiform Gyrus | 0.105 | 0.118 | 0.088 | 0.086 |
| Left Occipital Pole | -0.212 | -0.082 | 0.070 | -0.203 |
| Left Paracingulate Gyrus | -0.109 | -0.024 | -0.042 | 0.065 |
| Left Parahippocampal Gyrus, anterior division | 0.048 | 0.086 | -0.069 | -0.146 |
| Left Parahippocampal Gyrus, posterior division | -0.012 | 0.058 | 0.062 | -0.116 |
| Left Parietal Operculum Cortex | 0.151 | -0.030 | -0.036 | -0.101 |
| Left Planum Polare | 0.041 | 0.075 | -0.010 | 0.219 |
| Left Planum Temporale | 0.139 | 0.000 | -0.060 | 0.079 |
| Left Postcentral Gyrus | -0.015 | -0.118 | -0.082 | -0.060 |
| Left Precentral Gyrus | -0.131 | -0.007 | -0.045 | -0.060 |
| Left Precuneous Cortex | 0.059 | 0.007 | -0.004 | -0.101 |
| Left Subcallosal Cortex | -0.065 | -0.078 | -0.140 | -0.002 |

| | | | | |
|---|---|---|---|---|
| Left Superior Frontal Gyrus | -0.020 | -0.126 | -0.050 | 0.026 |
| Left Superior Parietal Lobule | -0.075 | -0.023 | 0.028 | -0.054 |
| Left Superior Temporal Gyrus, anterior division | 0.052 | 0.140 | -0.072 | 0.148 |
| Left Superior Temporal Gyrus, posterior division | -0.150 | 0.047 | 0.002 | -0.093 |
| Left Supracalcarine Cortex | -0.004 | -0.077 | 0.181 | -0.093 |
| Left Supramarginal Gyrus, anterior division | 0.029 | 0.005 | -0.033 | 0.114 |
| Left Supramarginal Gyrus, posterior division | 0.167 | 0.100 | 0.090 | -0.027 |
| Left Temporal Fusiform Cortex, anterior division | 0.045 | 0.076 | -0.059 | 0.062 |
| Left Temporal Fusiform Cortex, posterior division | -0.052 | 0.124 | 0.069 | 0.182 |
| Left Temporal Occipital Fusiform Cortex | 0.089 | -0.049 | -0.054 | -0.100 |
| Left Temporal Pole | 0.015 | 0.073 | 0.025 | 0.187 |
| Right Angular Gyrus | -0.015 | -0.032 | 0.063 | -0.074 |
| Right Central Opercular Cortex | -0.120 | -0.020 | -0.063 | -0.070 |
| Right Cingulate Gyrus, anterior division | 0.027 | 0.176 | 0.038 | 0.036 |
| Right Cingulate Gyrus, posterior division | 0.092 | -0.058 | 0.051 | -0.256 |
| Right Cuneal Cortex | -0.111 | -0.174 | 0.015 | -0.224 |
| Right Frontal Medial Cortex | -0.119 | -0.072 | -0.060 | -0.011 |
| Right Frontal Operculum Cortex | -0.032 | -0.015 | -0.019 | -0.056 |
| Right Frontal Orbital Cortex | -0.018 | -0.040 | -0.104 | 0.076 |
| Right Frontal Pole | -0.018 | -0.050 | -0.095 | 0.058 |
| Right Heschl's Gyrus (includes H1 and H2) | -0.025 | -0.084 | -0.079 | -0.138 |
| Right Inferior Frontal Gyrus, pars opercularis | -0.102 | -0.050 | -0.156 | -0.094 |
| Right Inferior Frontal Gyrus, pars triangularis | -0.215 | -0.062 | -0.185 | -0.128 |
| Right Inferior Temporal Gyrus, anterior division | 0.038 | -0.076 | -0.097 | -0.180 |
| Right Inferior Temporal Gyrus, posterior division | 0.069 | -0.002 | -0.071 | -0.128 |
| Right Inferior Temporal Gyrus, temporooccipital part | -0.153 | -0.115 | -0.171 | -0.217 |
| Right Insular Cortex | -0.035 | 0.031 | -0.006 | -0.068 |
| Right Intracalcarine Cortex | -0.078 | -0.140 | -0.117 | -0.088 |
| Right Juxtapositional Lobule Cortex (formerly Supplementary Motor Cortex) | -0.013 | -0.167 | -0.114 | -0.135 |
| Right Lateral Occipital Cortex, inferior division | -0.055 | -0.088 | 0.125 | 0.008 |
| Right Lateral Occipital Cortex, superior division | -0.035 | -0.078 | 0.043 | -0.119 |
| Right Lingual Gyrus | -0.180 | -0.115 | -0.096 | -0.149 |
| Right Middle Frontal Gyrus | 0.117 | 0.014 | 0.102 | 0.055 |
| Right Middle Temporal Gyrus, anterior division | -0.130 | -0.068 | -0.120 | -0.126 |
| Right Middle Temporal Gyrus, posterior division | -0.030 | -0.033 | -0.008 | 0.045 |
| Right Middle Temporal Gyrus, temporooccipital part | -0.008 | -0.071 | 0.010 | -0.109 |
| Right Occipital Fusiform Gyrus | 0.115 | 0.249 | 0.163 | 0.237 |
| Right Occipital Pole | -0.053 | -0.067 | 0.074 | -0.006 |
| Right Paracingulate Gyrus | -0.062 | 0.123 | 0.042 | -0.085 |
| Right Parahippocampal Gyrus, anterior division | -0.093 | -0.061 | -0.033 | -0.109 |
| Right Parahippocampal Gyrus, posterior division | -0.105 | -0.118 | -0.103 | -0.137 |

| | | | | |
|---|---|---|---|---|
| Right Parietal Operculum Cortex | 0.105 | -0.050 | -0.019 | -0.072 |
| Right Planum Polare | -0.033 | 0.050 | -0.042 | -0.148 |
| Right Planum Temporale | -0.064 | -0.135 | -0.057 | -0.209 |
| Right Postcentral Gyrus | -0.071 | 0.014 | 0.047 | 0.051 |
| Right Precentral Gyrus | 0.031 | 0.027 | -0.039 | -0.175 |
| Right Precuneous Cortex | 0.077 | 0.071 | 0.010 | -0.035 |
| Right Subcallosal Cortex | 0.018 | 0.044 | -0.007 | -0.106 |
| Right Superior Frontal Gyrus | -0.159 | -0.162 | -0.149 | -0.156 |
| Right Superior Parietal Lobule | -0.007 | 0.035 | -0.001 | -0.063 |
| Right Superior Temporal Gyrus, anterior division | -0.155 | -0.045 | -0.124 | -0.175 |
| Right Superior Temporal Gyrus, posterior division | -0.133 | -0.052 | -0.127 | -0.081 |
| Right Supracalcarine Cortex | -0.131 | -0.153 | 0.138 | -0.174 |
| Right Supramarginal Gyrus, anterior division | -0.101 | -0.061 | -0.078 | -0.068 |
| Right Supramarginal Gyrus, posterior division | -0.149 | -0.079 | -0.022 | -0.222 |
| Right Temporal Fusiform Cortex, anterior division | -0.005 | 0.064 | -0.020 | -0.041 |
| Right Temporal Fusiform Cortex, posterior division | -0.095 | -0.036 | 0.030 | -0.137 |
| Right Temporal Occipital Fusiform Cortex | -0.124 | -0.080 | 0.002 | -0.073 |
| Right Temporal Pole | -0.181 | -0.112 | -0.230 | -0.284 |

*$p_{FDR} < 0.05$; **$p_{FDR}$*

**Supplementary Table 9.** Contributions of selected region of interests (ROIs) to each principle components (PCs) resulted from of the Singular Value Decomposition (SVD) analyses for $SD_{BOLD}$, $SD_{DELTA}$, and $SD_{THETA}$. ROIs derived from the previous fMRI literature, and their corresponding ROIs in Harvard-Oxford atlas to investigate the age-dependent relationship between TMT and $SD_{BOLD}$ or $SD_{EEG}$ (See: Table 1). As a criterion, the minimum total variance explained over 70% was selected (Jollife and Cadima, 2016), resulting in three PCs in $SD_{BOLD}$ (52.82%, 10.34%, and 7%), two PCs in $SD_{DELTA}$ (67.37%, 10.95%), and one PC in $SD_{THETA}$ (75.63%).

| | **$SD_{BOLD}$** | | | **$SD_{DELTA}$** | | **$SD_{THETA}$** |
|---|---|---|---|---|---|---|
| **Regions of Interest** | PC1 | PC2 | PC3 | PC1 | PC2 | PC1 |
| Left.Cingulate.Gyrus..anterior.division | 6.089654 | 10.83573301 | 0.57335642 | 6.293254 | 4.771 | 6.074438 |
| Left.Cingulate.Gyrus..posterior.division | 5.157258 | 10.43198257 | 14.34970502 | 5.512103 | 8.818 | 5.977285 |
| Left.Frontal.Medial.Cortex | 2.291563 | 7.93654130 | 10.93536588 | 3.528656 | 0.025 | 4.072425 |
| Left.Insular.Cortex | 6.077573 | 2.00659998 | 14.77254088 | 6.299486 | 2.866 | 6.054641 |
| Left.Middle.Frontal.Gyrus | 7.236614 | 1.52487430 | 0.87793136 | 6.234210 | 0.0249 | 6.137876 |
| Left.Middle.Temporal.Gyrus..anterior.division | 3.086762 | 17.51744070 | 6.19688362 | 5.172188 | 15.0212 | 5.003268 |
| Left.Middle.Temporal.Gyrus..posterior.division | 4.406356 | 15.17712851 | 3.05589786 | 5.799520 | 10.474 | 5.435104 |
| Left.Middle.Temporal.Gyrus..temporooccipital.part | 5.472472 | 0.04546836 | 2.61039871 | 4.196545 | 1.262 | 4.698550 |
| Left.Precentral.Gyrus | 6.217582 | 2.57399365 | 4.67714091 | 6.894773 | 0.0000 | 6.686272 |
| Left.Superior.Frontal.Gyrus | 7.381836 | 0.32612274 | 0.61245518 | 5.792227 | 5.0748 | 5.692948 |
| Left.Superior.Temporal.Gyrus..anterior.division | 4.947440 | 0.05546047 | 0.03483502 | 4.491588 | 18.9260 | 4.466424 |
| Left.Superior.Temporal.Gyrus..posterior.division | 5.338529 | 7.26165079 | 0.67087115 | 5.690691 | 12.248 | 5.451148 |
| Right.Cingulate.Gyrus..anterior.division | 6.152739 | 9.90571267 | 0.90070689 | 6.200047 | 5.448 | 6.070030 |
| Right.Cingulate.Gyrus..posterior.division | 5.192974 | 10.04779651 | 14.34718789 | 5.404548 | 9.346 | 5.913883 |
| Right.Insular.Cortex | 7.339789 | 0.25286095 | 0.58469394 | 6.523552 | 0.0871 | 6.381107 |
| Right.Middle.Frontal.Gyrus | 5.606567 | 0.89429285 | 1.19883860 | 5.608391 | 1.1479 | 5.755169 |
| Right.Occipital.Fusiform.Gyrus | 5.520859 | 1.04779909 | 16.02672218 | 3.929315 | 2.298 | 3.899637 |
| Right.Precentral.Gyrus | 6.483434 | 2.15854154 | 7.57446847 | 6.428907 | 2.1581 | 6.229797 |

**Supplementary Table 10.** Results of multiple linear regression analyses investigating the relationship between the principle components (PCs) in $SD_{BOLD}$, $SD_{DELTA}$, and $SD_{THETA}$ derived from Singular Value Decomposition (SVD) and task completion time in Trail Making Test (Reitan, 1955; Reitan and Wolfson, 1995) as the dependent variable.

| | **$SD_{BOLD}$** | | | | **$SD_{DELTA}$** | | | | **$SD_{THETA}$** | | | |
|---|---|---|---|---|---|---|---|---|---|---|---|---|
| | **TMT-A** | | **TMT-B** | | **TMT-A** | | **TMT-B** | | **TMT-A** | | **TMT-B** | |
| | Beta | p-value | Beta | p-value | Beta | p-value | Beta | p-value | Beta | p-value | Beta | p-value |
| PC1 | 0.004 | 0.254 | 0.004 | 0.922 | 0.005 | 1 | 0.006 | 1 | -0.007 | 0.804 | -0.007 | <0.001 |
| Age | 0.009 | <0.001 | 0.015 | <0.001 | 0.012 | <0.001 | 0.009 | <0.001 | 0.008 | <0.001 | 0.01 | <0.001 |
| PC1 * Age | 0.0002 | 0.592 | 0.0006 | 0.961 | 0.0001 | 0.481 | -0.0002 | 0.325 | 0.0002 | 0.372 | -0.0002 | 0.161 |
| PC2 | -0.017 | 0.961 | -0.017 | 0.186 | -0.011 | 1 | 0.003 | 0.764 | | | | |
| PC2 * Age | -0.0009 | 0.296 | -0.002 | **0.027** | -0.0005 | 0.362 | -0.001 | 0.08 | | | | |
| PC3 | 0.02 | 0.142 | 0.019 | 0.282 | | | | | | | | |
| PC3 * Age | -0.0002 | 0.824 | 0.0003 | 0.556 | | | | | | | | |
